# Learning molecular determinants of selective small-molecule partitioning across biomolecular condensates

**DOI:** 10.64898/2026.05.23.727301

**Authors:** Aahil Khambhawala, Shiv Rekhi, Qizan Chen, Priyesh Mohanty, Daniel P. Tabor, Jeetain Mittal

## Abstract

The functional role of biomolecular condensates is shaped by the composition of constituent proteins, nucleic acids, ions, and small molecules. Selective partitioning of small molecules into condensates has therefore emerged as a potential route to condensate-specific chemical probes and therapeutics. Although partitioning is influenced by differences in solvation environments between coexisting dense and dilute phases, a molecular framework connecting small-molecule structure to condensate-specific enrichment remains lacking. Here, we use existing experimental partitioning data for a library of FDA-approved drugs and metabolites across four biomolecular condensates to develop an interpretable graph-based model of small-molecule partitioning. By combining multitask pretraining, condensate-specific fine-tuning, evidential uncertainty quantification, and atom-level attribution analysis, our model predicts continuous partition coefficients with improved accuracy over descriptor-based approaches. Atom-level attributions reveal that condensate partitioning is not governed by a universal chemical rule: the same molecular scaffold can be read differently by distinct condensate environments, with local atomic context and connectivity determining whether specific atoms promote or suppress enrichment. We further apply the trained model to ~1.7 million drug-like molecules from ChEMBL, identifying a chemically diverse space of predicted condensate-selective partitioners and mapping regions where predictions are confident versus where new measurements would be most informative. Together, this work establishes condensate partitioning as a chemically learnable property shaped by the interplay between small-molecule structure and condensate-specific microenvironments, providing an interpretable and uncertainty-aware framework for defining molecular determinants of partitioning and guiding the discovery of condensate-selective small molecules.

## Introduction

The phase separation of biomolecules into biomolecular condensates creates dense phases with chemical environments distinct from the surrounding dilute phase. These environments are critical for condensate function [1–4]. It is increasingly clear that aberrant condensation contributes to disease, notably neurodegeneration and cancer [5–9], leading to interest in targeting condensates for therapeutic intervention [7,10,11]. Small molecules can modulate the thermodynamics and dynamics of condensates, as well as regulate dense-phase composition, all of which are critical for their physiological function [12,13]. Thus, there is significant interest in the design and development of condensate-modulating small molecules.

A major hurdle in developing condensate-modulating molecules is specificity: a useful molecule should preferentially act on the target condensate while minimizing effects on other biomolecular assemblies [14–17]. Klein et al [18] first demonstrated that cancer therapeutics can partition selectively into nuclear condensates, influencing drug pharmacodynamics and contributing to cancer resistance. This observation raised the possibility that condensates present chemically distinct environments that can differentially enrich small molecules. However, the molecular features that determine whether a compound partitions into one condensate, many condensates, or none remain incompletely understood.

Subsequent studies have expanded this observation from cancer therapeutics to larger small-molecule libraries. Kilgore et al [19] studied the partitioning of more than 1500 fluorescent probes into four intrinsically disordered polypeptide (IDP) condensates and observed condensate-dependent enrichment profiles. Using graph-neural network-based structure-activity relationships, they showed that molecular structure contains information predictive of binary partitioning behavior. Thody et al.[20] quantified the partitioning of approximately 1700 FDA-approved drugs and metabolites into multidomain protein condensates using mass spectrometry. In that dataset, partition coefficients (PCs) spanned nearly six orders of magnitude, and descriptor-based models identified features related to solubility and hydrophobicity as important predictors of enrichment. Complementary simulations and sequence-based approaches have connected small-molecule enrichment to condensate micropolarity, dense-phase composition, and the interaction networks formed by the molecules within the condensate [18,20,21]. Together, these studies establish that small-molecule partitioning reflects both molecular properties of the compound and the chemical environment of the condensate. However, an atom-level framework for predicting continuous partitioning strength, explaining condensate selectivity, or identifying where model predictions are reliable remains lacking.

Despite these advances, detailed examination of the experimental data reveals systematic deviations from simple physicochemical trends. Molecules with nearly identical hydrophobicity (as measured by water/octanol partition coefficients) or high structural similarity can display markedly different partition coefficients, as illustrated in **Fig. 1a,b**. In some cases, small changes in functional group identity, heteroatom placement, or molecular connectivity are associated with large differences in condensate enrichment. Conversely, molecules with similar water/octanol partition coefficients (logP) can show substantially different dilute-to-dense phase partition coefficients (logPC) (**Fig. 1c**). These observations suggest that bulk hydrophobicity and global structural similarity alone are insufficient to explain condensate partitioning across chemical space. Instead, condensate selectivity likely depends on local atomic environments, molecular connectivity, and higher-order structural context in addition to bulk solvation properties.

**Fig 1.**
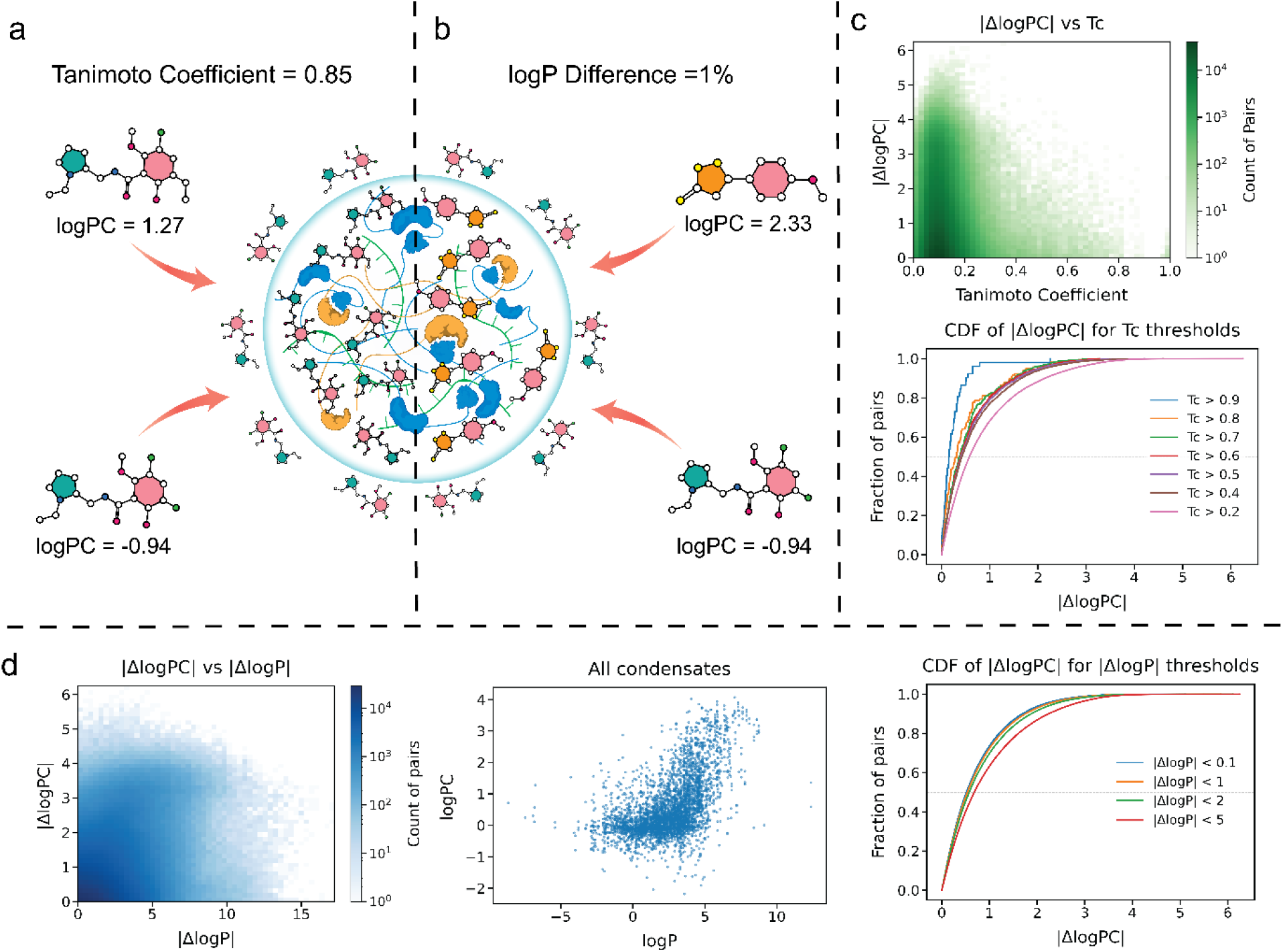
Structural similarity and hydrophobicity alone do not reliably predict condensate partitioning. (a) Schematic illustrating that molecules with high structural similarity, quantified by Tanimoto coefficient, can exhibit substantially different partition coefficients (logPC). (b) Example showing that molecules with nearly identical hydrophobicity, measured by logP, can also display large differences in logPC. (c) Dataset-wide analysis of differences in condensate partitioning as a function of molecular similarity. The upper panel shows the density of molecular pairs binned by Tanimoto coefficient and absolute logPC difference, and the lower panel shows cumulative distribution functions of |ΔlogPC| for different Tanimoto similarity thresholds. (d) Dataset-wide analysis of differences in condensate partitioning as a function of hydrophobicity similarity. The left panel shows pairwise density as a function of |ΔlogP| and |ΔlogPC|, the middle panel shows overall logPC versus logP across condensates, and the right panel shows cumulative distributions of |ΔlogPC| for different |ΔlogP| thresholds. Together, these analyses show that global structural similarity and bulk hydrophobicity capture broad trends but are insufficient to explain condensate-specific partitioning across chemical space.

In this work, we develop an interpretable graph-based framework to identify atomic determinants of small-molecule partitioning into biomolecular condensates. The model combines multitask pretraining on solvation-related molecular properties, condensate-specific fine-tuning on experimental partitioning data, and evidential uncertainty quantification [22]. This framework predicts continuous logPC values across four condensates with improved performance relative to descriptor-based models, while providing uncertainty estimates for each prediction. Using SHapley Additive exPlanations (SHAP) [23], we quantify atom-level contributions to predicted partitioning and identify how the same molecular scaffold can be interpreted differently by distinct condensate environments. Finally, we apply the trained model to approximately 1.7 million drug-like molecules from ChEMBL to map predicted condensate-selective chemical space and identify regions where new measurements would be most informative [24]. Together, these results provide an atom-level and uncertainty-aware framework for connecting molecular structure to condensate-specific partitioning [25].

## Results & Discussion

### Molecular graphs improve generalization in predicting condensate partitioning

Descriptor-based models rely on user-defined features in combination with classical machine learning approaches to predict partitioning. Given the limited experimental data available for condensate partitioning and the prior success of descriptor-based models in predicting logPC values, we trained XGBoost [26] and LightGBM [27] models for all four condensates using two descriptor sets QikProp descriptors and RDKit two-dimensional and three-dimensional molecular descriptors. To evaluate generalization beyond closely related molecules, we constructed the train, validation, and test splits such that they share limited structural similarity and restricted scaffold overlap (**Fig. S1-S2**) Across the four condensates, models trained on two-dimensional QikProp features recovered a substantial fraction of the variation in experimental logPC values in the training set. Additionally, t-SNE visualizations generated for each condensate showed that inclusion of three-dimensional molecular descriptors increased the apparent separation between compounds with distinct partitioning behavior (**Figs. S3-S10**). **Table S1** further supports this observation quantitatively, with separability metrics for these various feature sets. Therefore, we extended the feature space to include three-dimensional descriptors derived from RDKit [28] and Mordred [29] further improved training-set performance and yielded tight parity plots for the training data (**Fig. S11**). However, on the test set, all descriptor-based models exhibited a clear gap between training and test performance (**Fig. S11**). This trend is evident across the full set of baseline comparisons in Figs. S3–S11: descriptor-based models often capture the training data well, but their predictions become less reliable for structurally separated test molecules, indicating limited transferability beyond closely related chemotypes. Combining the XGBoost and LightGBM models in a stacked ensemble using kernel and ridge regressors yielded only modest improvements over the best individual models on the test set and did not remove these generalization errors. **Table S2** shows the detailed hyperparameter values for the best model for all 4 condensates Next, we tested whether a model that learns directly from molecular graph structure, rather than relying on predefined molecular descriptors, could improve prediction on the test set. We used a message passing graph-neural network [30,31], with node and edge features that encode the chemistry of the molecule (**see Methods**). Because graph-based models often require larger datasets to realize their representation learning advantages [32], we pretrained the model [33–35] on logP, logS, and CIlogS values for 150,000 small molecules (**Fig. 2**). These properties were selected because they report on molecular solvation and hydrophobicity, which are expected to contribute to condensate partitioning while not fully determining condensate-specific enrichment. We anticipated that pretraining would allow the GNN to learn molecular features associated with solvation, and the subsequent fine-tuning on condensate partitioning data would adapt these features to account for the distinct chemical environments of biomolecular condensates.

**Fig. 2.**
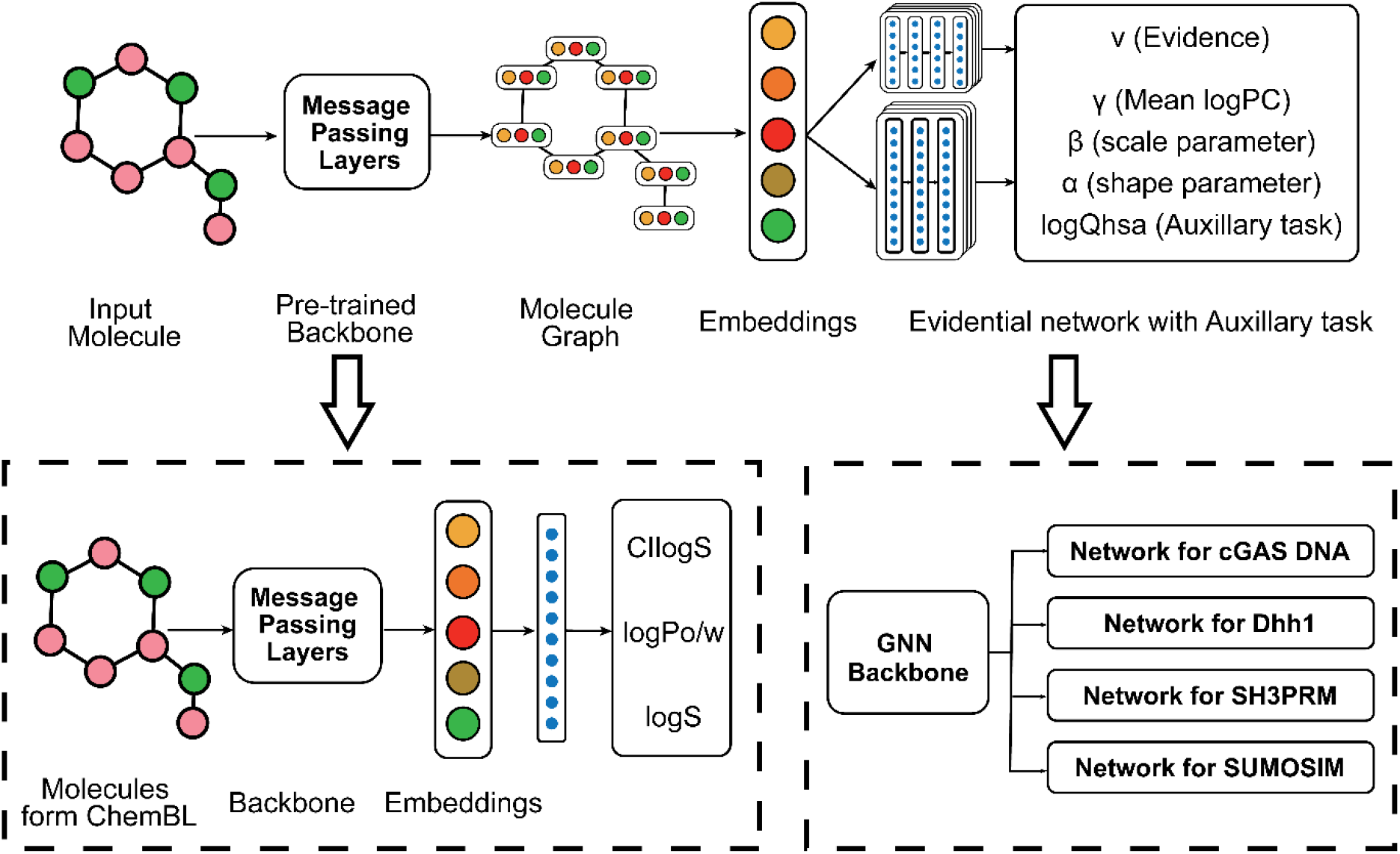
Graph neural network framework for condensate-specific partitioning prediction with evidential uncertainty. The top panel illustrates the overall pipeline. An input molecule is represented as a molecular graph and processed by a pretrained message passing backbone to generate molecular embeddings. These embeddings are passed to condernsate-specific evidential regression heads that output the predicted mean logPC and Normal-Inverse-Gamma parameters used to estimate predictive uncertainty, along with an auxiliary logQhsa prediction. The bottom left panel shows pretraining of the backbone on ChemBL molecules using solvation-related physicochemical properties, including CIlogS, logP, and logS. The bottom right panel shows fine-tuning of the shared GNN backbone into separate condensate-specific networks for cGAS DNA, Dhh1, SH3PRM, and SUMOSIM.

The pretraining step produced a model that accurately predicted all three solvation-related properties, with train and test metrics reported in **Table S3**. Using this model as a backbone, we then fine-tuned the network to learn condensate-specific partitioning. We trained the model on molecules for which logPC values were available for all four condensate systems. Downstream of the shared pretrained backbone, we used four independent task-specific heads to predict logPC values for each condensate. We also selected logQhsa as an auxiliary task because it was moderately correlated with condensate partitioning, providing a related physicochemical signal while still allowing the model to capture deviations specific to condensate partitioning. Detailed comparisons of model performance without pretraining, without the auxiliary task and without the multitask architecture are provided in **Figs. S12-S15** and **Table S4**.

On the held-out test set, the multitask evidential GNN showed improved performance relative to descriptor-based models across all four condensates (**Fig. 3, Table 1)**. Training-set performance with uncertainty bars is shown in **Fig. S16**. To assess model robustness, we trained additional models with backbone sizes larger and smaller than the current configuration. Across fifty random initializations, the present model exhibited the most stable and optimal performance, balancing predictive accuracy and uncertainty calibration (**Fig. S17**). Among the four systems, the model achieved the lowest MAE for Dhh1, followed by SH3PRM and cGAS DNA, whereas SUMOSIM showed a smaller improvement relative to the descriptor-based model (**Table 1** Overall, these results demonstrate that a pretrained evidential multitask GNN improves prediction of condensate partitioning relative to descriptor-based models for the training size and scaffold-aware splits used here. Having established this improved generalization, we next asked whether the learned graph representation could reveal atomic features associated with condensate-specific partitioning.

**Fig. 3.**
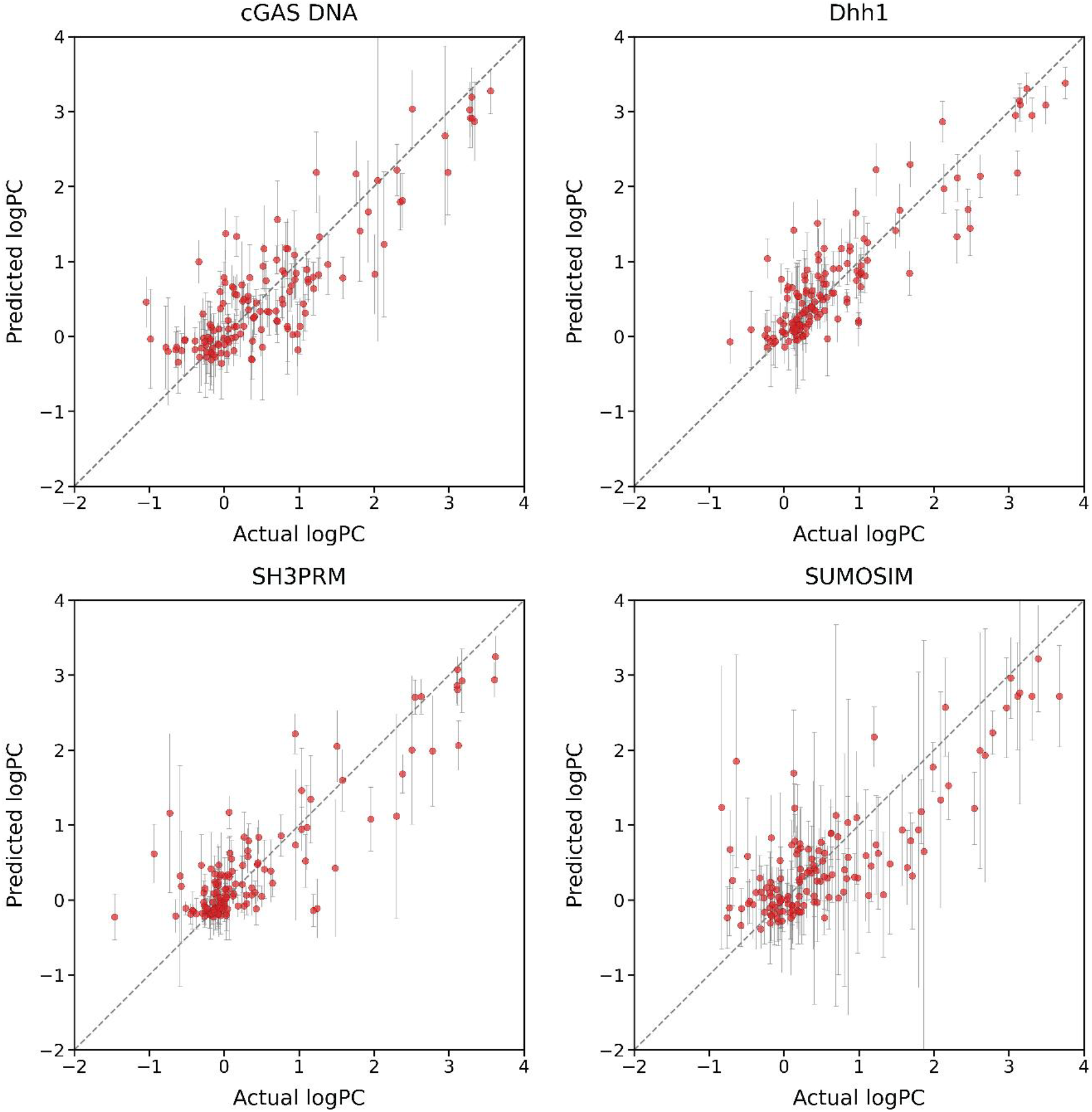
Predicted and experimental partition coefficients for held-out test molecules across four condensates. Parity plots compare predicted and experimental logPC values for cGAS DNA, Dhh1, SH3PRM, and SUMOSIM. Each point represents one molecule in the held-out test set, the dashed line denotes ideal agreement, and vertical error bars indicate predictive uncertainty estimated by the evidential regression model. The model captures condensate-dependent variation in logPC across structurally separates test molecules, with performance metrics summarized in Table 1.

**Table 1:** Comparison of descriptor-based baseline and graph neural network model performance on held-out test sets.

| Condensate | Baseline Model |  | GNN Model |  |
| --- | --- | --- | --- | --- |
|  | Test R<sup>2</sup> | Test MAE | Test R<sup>2</sup> | Test MAE |
| cGAS DNA | 0.59 | 0.45 | 0.75 | 0.38 |
| Dhh1 | 0.63 | 0.45 | 0.80 | 0.28 |
| SH3PRM | 0.62 | 0.44 | 0.77 | 0.30 |
| SUMOSIM | 0.56 | 0.46 | 0.62 | 0.44 |

### Atom-Level Attributions Reveal Condensate-Specific Drivers of Partitioning

We use SHapley Additive exPlanations (SHAP) analysis applied to individual atoms of the molecules to understand the atomic drivers of small-molecule partitioning [36,37]. SHAP analysis quantifies how much each atom within a molecule contributes to its predicted logPC by selectively “masking” or zeroing out atomic features while preserving the rest of the graph (**see Methods**). This approach allows us to ask whether the model relies on the same molecular features across condensates or whether distinct condensate environments assign importance to different atoms within the same molecule. To enable comparison between molecules of different sizes on a common scale, we normalize the SHAP values by the additive contributions of all atoms within the molecule. SHAP values close to 0 indicate that masking those atoms does not significantly alter the predicted logPC, while negative or positive values indicate that the atoms reduce or increase the predicted logPC, respectively.

We first used representative molecule-level attribution maps to determine whether similar overall partitioning predictions arise from similar or distinct atomic contributions across condensates. For this, we selected two representative molecules with similar predicted logPC values across all four condensates (**Fig. 4a**). Despite their comparable overall partitioning behavior, the atom-level attribution maps differ substantially between condensates. For the first molecule, which exhibits weak partitioning across all four systems (top row, logPC ranging from −0.64 to −0.74), different condensate models assign the largest contributions to different regions of the molecule, including heterocyclic atoms, oxygen-containing linkages, and aromatic groups. For the second molecule, which partitions favorably into all four condensates (bottom row, logPC ranging from 2.26 to 2.79), the models again emphasize different atomic regions, with positive contributions distributed across distinct aromatic, heteroatomic, and linker environments dependent on condensate identity. exhibits even more pronounced condensate-specific attribution patterns. This molecule shows favorable partitioning into all four condensates, yet the atomic basis for these predictions varies dramatically. Thus, similar predicted logPC values can arise from different atomic attribution patterns, suggesting that each condensate model uses a distinct molecular context to predict enrichment.

**Fig. 4.**
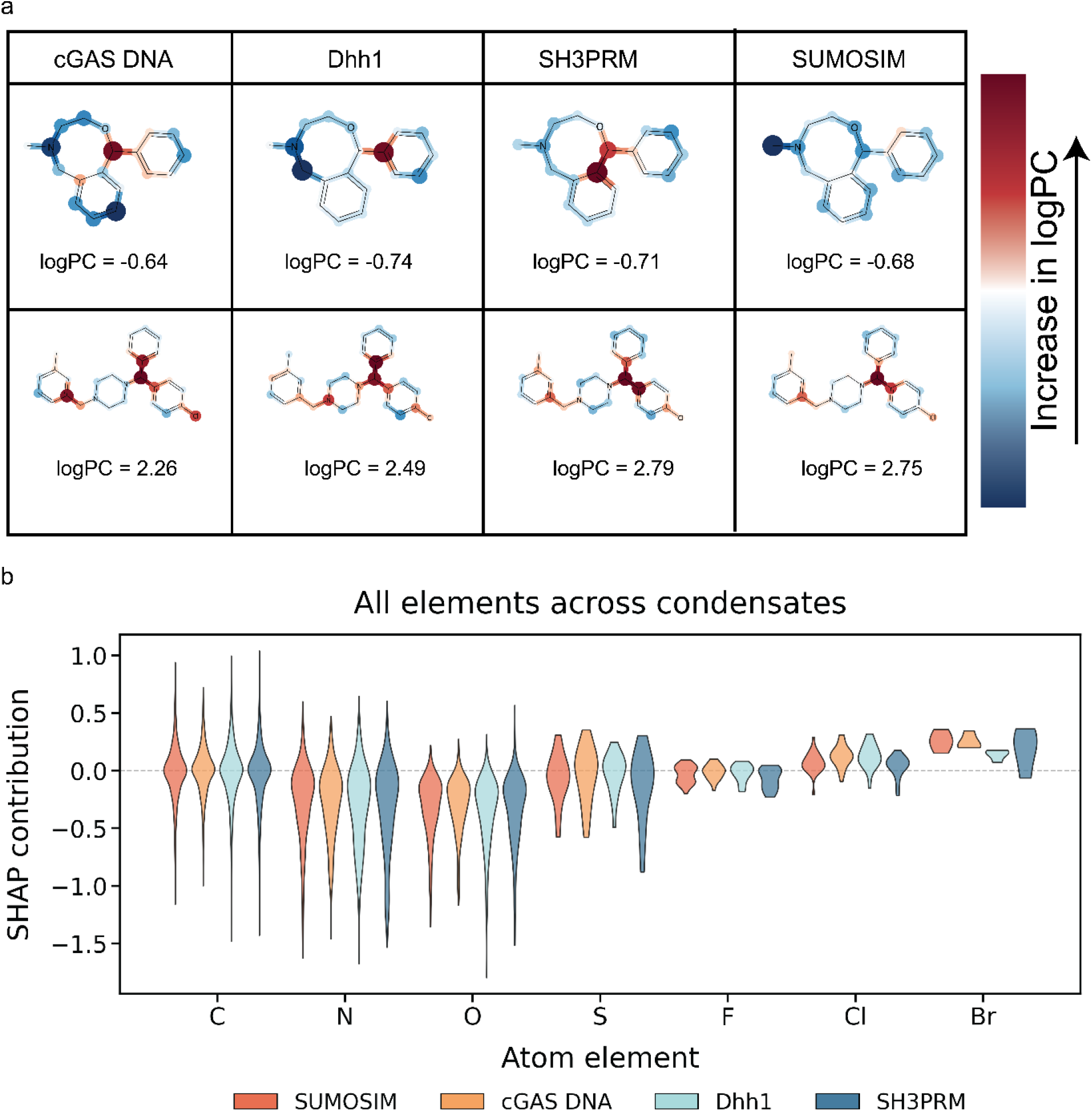
Atom-level attributions reveal condensate-specific drivers of small molecule partitioning. (a) Atom-level SHAP maps for two representative molecules evaluated across cGAS DNA, Dhh1, SH3PRM, and SUMOSIM condensates. The top row shows a molecule with weak predicted partitioning, and the bottom row shows a molecule with favorable predicted partitioning across the four condensates. Atoms are colored by their normalized SHAP contribution to the predicted logPC, with positive values indicating atoms that promote partitioning and negative values indicating atoms that suppress partitioning. Although the overall logPC values are similar across condensates for each molecule, the spatial distribution and magnitude of atomic SHAP contributions differ between condensates, indicating context-dependent atomic drivers of partitioning. (b) Violin plots of SHAP contributions grouped by atom element and condensate. Each distribution reflects atom-wise contributions to predicted logPC across the dataset. Broad and overlapping distributions across element types indicate that atomic identity alone does not determine partitioning; instead, atomic contributions depend on local molecular context and condensate identity.

These examples indicate that the learned models do not apply a single universal partitioning rule across condensates. Instead, the same chemical scaffold can be weighed differently depending on the biomolecular environment. We interpret these differences as evidence that condensate partitioning depends not only on global molecular properties, such as hydrophobicity, but also on the local atomic context and connectivity through which those properties are expressed. Such context dependence is consistent with the expectation of chemically distinct environments presented by nucleic-acid-containing condensates, such as cGAS DNA and Dhh1, and protein-only condensates, such as SH3PRM and SUMOSIM [38].

To determine whether these molecule-specific examples reflect broader trends across the dataset, we next aggregated SHAP contributions by atomic element type (**Fig. 4b**). Across all condensates, most elements display broad distributions spanning both positive and negative SHAP contributions, indicating that their influence on partitioning is highly context-dependent rather than uniformly favorable or unfavorable. Carbon and nitrogen atoms show the widest range of contributions, demonstrating that these common elements can either promote or suppress partitioning depending on their position within the molecular structure and local chemical environment. Oxygen atoms similarly exhibit substantial variability, with contributions distributed broadly around zero across all four condensates. Sulfur and halogen atoms show more constrained distributions, although their contributions also depend on molecular context. Among the halogens, the distributions shift toward more positive contributions from F to Cl and Br, suggesting that larger and more polarizable substituents may more often favor predicted partitioning, although the overlap between distributions indicates that this trend remains context dependent.

Importantly, the substantial overlap in element-level distributions across condensates indicates that no single element consistently acts as a universal driver or suppressor of partitioning. Instead, atomic contributions depend on both the structural context of the atom within the molecule and the condensate environment being modeled. Together, the molecule-level attribution maps and dataset-wide element analysis support a picture in which condensate partitioning is governed by context-dependent molecular recognition than simple element-based or hydrophobicity-based rules.

### Virtual Screening Maps Condensate-Selective Chemical Space

Given the improved performance of the models trained in this work, we sought to characterize the chemical landscape of predicted condensate partitioning across drug-like molecules. We applied the evidential graph neural networks to approximately 1.72 million drug-like compounds from the ChEMBL database [39] and predicted their partitioning into the four condensate systems.

Across the ChEMBL library, predicted logPC values span a broad range for all four condensates (**Fig. 5a**). Most molecules fall outside the strongly enriched regime, with many predicted to be excluded or only weakly enriched in the dense phase. Nevertheless, each condensate exhibits a substantial high-partitioning tail, with approximately 30% of screened molecules predicted to have logPC > 1 in at least one condensate. This corresponds to more than 500,000 candidate molecules with strong predicted condensate enrichment, indicating that condensate partitioning potential is broadly represented across drug-like chemical space than restricted to a narrow class of scaffolds. The four condensate systems display broadly similar distribution shapes, consistent with shared contributions from general physicochemical properties, including hydrophobicity and solvation. However, differences between condensates are most informative in the intermediate logPC regime, where molecules are neither strongly excluded nor strongly enriched across all condensates and where condensate-specific interactions are most likely to influence predicted selectivity. Because these predictions extend beyond the experimentally measured training set, we interpret this screen as a map of candidate chemical space rather than as direct evidence of experimentally validated partitioning.

**Fig 5.**
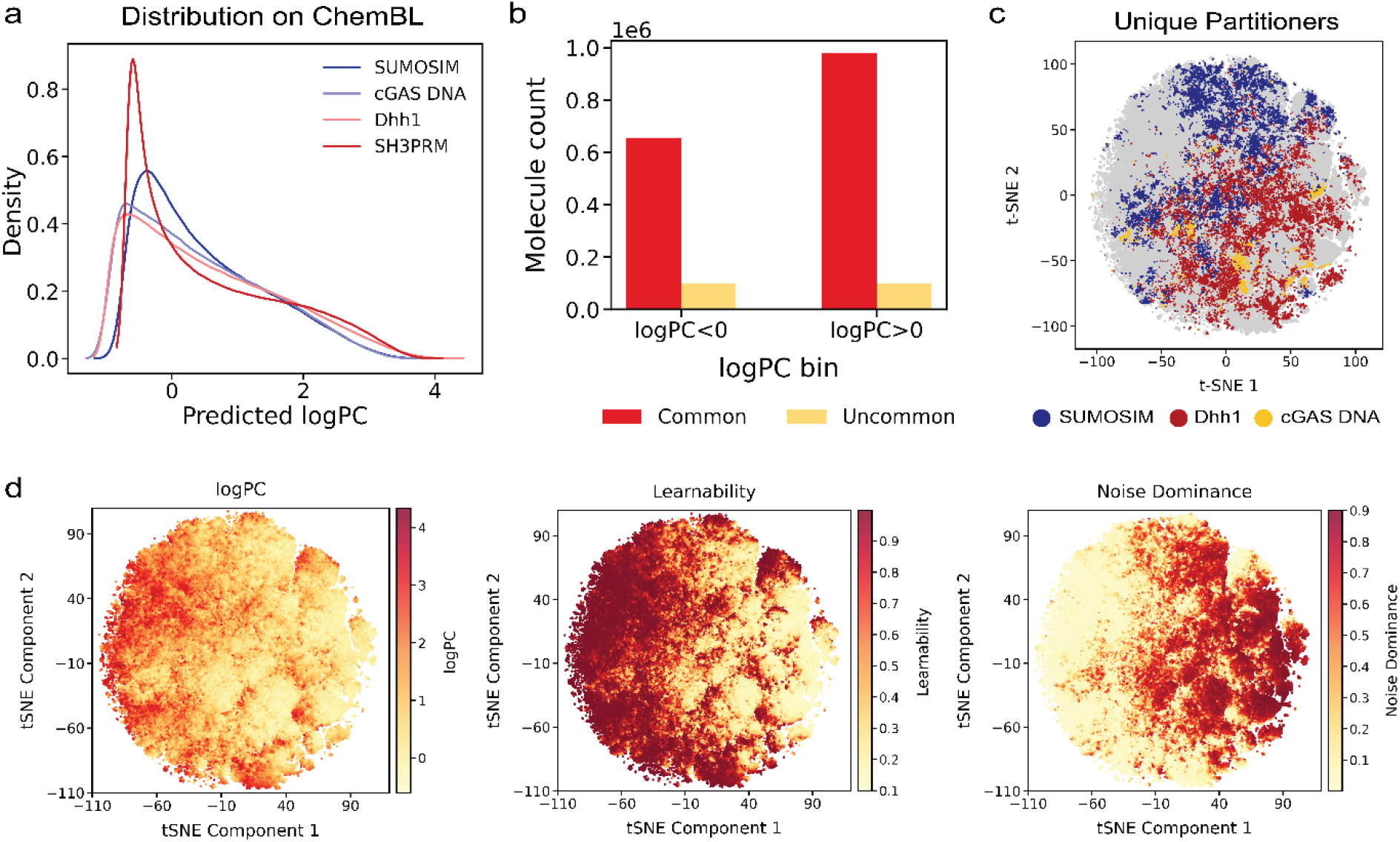
Large-scale virtual screening maps condensate partitioning across drug-like chemical space. (a) Distribution of predicted logPC values for approximately 1.72 million ChEMBL molecules across cGAS DNA, Dhh1, SH3PRM, and SUMOSIM condensates. (b) Counts of molecules classified as common or selective based on predicted partitioning behavior across all four condensates. Common molecules show the same qualitative behavior across all condensates, whereas selective molecules are predicted to partition into only a subset of condensates. (c) t-SNE projection of learned molecular embeddings highlighting predicted unique partitioners, defined as molecules predicted to partition into one condensate while being excluded from the other three. (d) t-SNE maps of the same chemical space colored by predicted logPC, learnability, and noise dominance derived from the evidential framework. Learnability highlights regions where additional data are expected to improve the model, whereas noise dominance highlights regions where uncertainty is more strongly associated with apparent data noise or variability.

To assess condensate selectivity, we categorized molecules according to their predicted partitioning patterns across all four condensates (**Fig. 5b**). Molecules were classified as common if they showed the same qualitative behavior across all four systems, either predicted enrichment in all condensates or predicted exclusion from all condensates. Molecules were classified as selective if they displayed different behavior, with predicted enrichment in some condensates and exclusion from others. Under this classification, common behavior was the dominant outcome: many molecules were predicted to be excluded from all condensates, while many positive partitioners were predicted to enrich in all four condensates. However, a substantial subset of molecules showed differential partitioning profiles, indicating that condensate-selective enrichment is accessible within drug-like chemical space. These predicted selective partitioners define candidate chemical matter for developing condensate-selective probes or modulators, although we emphasize that selectivity inferred from the screen should be viewed as a prioritization criterion for future experiments rather than as a validated property.

Motivated by this predicted differential partitioning, we next identified unique partitioners, defined as molecules with predicted favorable partitioning into only one condensate and predicted exclusion from the other three condensates. This analysis revealed marked differences across condensate systems. SUMOSIM had the largest population of predicted unique partitioners under this definition (16,441 molecules), followed by Dhh1 (10,860 molecules) and cGAS DNA (1,781 molecules), whereas SH3PRM showed no unique partitioners under this definition. The absence of predicted SH3PRM-unique partitioners does not imply that selective SH3PRM-targeting molecules cannot exist; rather, it indicates that such molecules were not identified within the screened ChEMBL chemical space and model-defined selectivity threshold.

To visualize where in chemical space the model makes confident predictions and where uncertainty dominates, we computed molecular representations using the learned graph embeddings from the trained evidential neural networks and projected them into two dimensions (**Fig. 5d**). Because these embeddings are high-dimensional, principal component analysis was first applied for compression, followed by t-SNE projection. The resulting maps show that predicted logPC values vary across chemical space and that high-partitioning molecules are distributed across multiple regions rather than confined to a single cluster together. This distribution suggests that strong predicted condensate partitioning can be achieved by diverse molecular scaffolds. Equivalent t-SNE maps for other condensates are shown in **Fig. S18**. An additional t-SNE map colored by logP shows a diffuse distribution of lipophilicity across the embedding space, suggesting that the logPC organization is not simply inherited from lipophilicity driven structure in the pretrained representation (**Fig. S19**). Additional analysis of molecules predicted to selectively partition into or be excluded from a single condensate shows that condensate specific behavior is associated with differences in molecular size, polarity, sp^3^ character, heteroatom content, and selected structural motifs (**Fig. S20**)

The evidential framework decomposes total predictive uncertainty into epistemic uncertainty, which reflects model uncertainty due to limited training data, and aleatoric uncertainty, which reflects irreducible noise or variability in the data. We define learnability as the ratio of epistemic uncertainty to total uncertainty and noise dominance as the ratio of aleatoric uncertainty to total uncertainty. The learnability map highlights regions where additional experimental measurements are expected to be most informative because uncertainty is dominated by limited model knowledge. In contrast, the noise-dominance map highlights regions where prediction uncertainty is more strongly associated with apparent data noise or assay variability, suggesting that improved measurement precision may be more valuable than simply increasing the number of related measurements.

Together, these uncertainty analyses provide a framework for prioritizing molecules for experimental validation and future model refinement. High-confidence candidates are those with strong predicted partitioning or selectivity, low overall uncertainty, and low noise dominance. Molecules in high-learnability regions, particularly those with predicted selectivity, represent valuable active-learning targets because their characterization would expand model coverage into underrepresented regions of chemical space. Thus, the ChEMBL screen serves as both a source of predicted condensate-selective candidates and an uncertainty-aware guide for future partitioning measurements.

## Conclusions and Outlook

In this work, we developed an interpretable and uncertainty-aware framework for predicting small-molecule partitioning into biomolecular condensates. By combining graph neural networks with evidential uncertainty quantification and atom-level attribution analysis, we connected molecular structure to condensate-specific partitioning behavior across four biomolecular condensates. The resulting model predicts continuous partition coefficients with improved performance relative to descriptor-based baselines, while also providing molecule-specific uncertainty estimates that can guide experimental prioritization.

Atomic-level SHAP analysis indicates that condensate partitioning is governed by context-dependent molecular recognition rather than universal chemical rules. Molecules with similar predicted logPC values across multiple condensates exhibit distinct attribution patterns, with different condensate models emphasizing different atomic regions within the same scaffold. Dataset-wide element-level analysis further shows that common atoms such as carbon, nitrogen, and oxygen can either promote or suppress predicted partitioning depending on their local molecular environment. These results support a picture in which condensate enrichment reflects the interplay between global molecular properties, local atomic context, and condensate identity.

Large scale virtual screening of approximately 1.72 million drug-like molecules from ChEMBL showed that predicted condensate partitioning is broadly distributed across drug-like chemical space. Although many molecules were predicted to behave similarly across condensates, a substantial subset displayed differential or unique partitioning profiles, suggesting that condensate-selective chemical matter is accessible within existing drug-like libraries. Because these predictions remain to be experimentally validated, we view the screen primarily as a prioritization resource: it identifies candidate molecules for testing and highlights regions of chemical space where new measurements would most efficiently improve model performance.

The absence of predicted unique partitioners for SH3PRM under our model-defined criteria suggests that some condensate environments may be more difficult to target selectively within the screened ChEMBL chemical space. However, this result should not be interpreted as evidence that SH3PRM-selective molecules cannot exist. Instead, it motivates future exploration of broader or more specialized chemical libraries, alternative selectivity thresholds, and experimental measurements designed to test model predictions. More generally, the uncertainty maps generated by the evidential framework provide a route to active learning, where molecules can be selected not only for high predicted partitioning or selectivity but also for their potential to improve model coverage. Future work should also test how partitioning rules learned from reconstituted condensates transfer to cellular environments, where competing condensates, biomolecular clients, condensate maturation, and cellular availability of small molecules may reshape apparent selectivity.

Together, this work establishes a foundation for rational discovery of condensate-selective small molecules by linking molecular structure, atom-level context, and macroscopic partitioning behavior. Future experimental validation of high-confidence and high-learnability candidates will be essential for converting these predictions into chemical probes and for refining the molecular grammar of condensate partitioning. More broadly, the framework developed here provides a general strategy for mapping how small molecules interact with the distinct, sequence-encoded environments of biomolecular condensates.

## Methodology

### Data Processing

The dataset used in this work was derived from Thody et al., who systematically quantified small-molecule partitioning across four representative condensate systems: SUMOSIM, cGAS DNA, Dhh1, and SH3PRM [20]. In their study, condensates were reconstituted in vitro using purified proteins or nucleic acid components, and the partitioning of each compound between the dense and dilute phases was measured using mass spectrometry under controlled buffer conditions. Each small molecule was incubated with preformed condensates, and partition coefficients were calculated from the ratio of signal intensities between the two phases. The resulting dataset provides experimentally determined partition coefficients across the four condensates.

From this dataset, only compounds with reported values for all four condensate systems were selected to ensure uniform supervision during multitask model development. The data were subsequently divided into training, validation, and test splits under strict criteria designed to prevent structural leakage. Cross-split duplicates were not allowed. Molecules with Tanimoto similarity greater than 0.6 to any training molecule were considered high-similarity cases and limited to less than twenty percent. Those exceeding 0.7 similarity were categorized as very high similarity and restricted to less than five percent. The proportion of mixed clusters, defined as molecular clusters spanning multiple partitions, was maintained below ten percent. Scaffold-level overlap, calculated using Jaccard similarity between Bemis-Murcko scaffolds, was kept below twenty percent to ensure minimal scaffold leakage. Graphs showing the data distribution in the training, validation, and test splits are provided in **Fig. S1**.

### Baseline Models

To provide a reference for model performance, we trained several baseline models on the same training, validation, and test splits. First, gradient boosted decision trees were implemented using XGBoost [26]. Second, an additional set of boosted tree models was trained using LightGBM [27]. Finally, we constructed a stacked ensemble model in which the predictions from the XGBoost and LightGBM models were combined using a kernel based regressor and a ridge regressor that learned optimal weights for the base model outputs. These baselines were evaluated with different molecular feature sets. Detailed description for the choice of feature sets is given in the Supporting Information. One variant used the two-dimensional QikProp features available in the data. A second variant used three-dimensional features generated using RDKit and Mordred [28,29]. A third variant combined all available features into a single input vector. Hyperparameters for each baseline model and for the stacked ensemble were optimized using Optuna [40]. Across these configurations, the baseline models fit the training data well but showed clear signs of overfitting, with substantially poorer generalization to test compounds.

### Graph neural Networks

Graph neural networks (GNNs) learn molecular representations directly from graph-structured inputs, where atoms are nodes and chemical bonds are edges [22,41]. Each atom is initially described by a feature vector encoding local physicochemical attributes, while bonds carry features capturing type or order. The network updates atomic representations through message-passing steps where each node iteratively exchanges information with its chemical neighborhood. Messages from neighboring atoms are aggregated and combined with current node states to produce updated embeddings. This process enables each atom to encode both intrinsic features and extended chemical context. After multiple propagation steps, node embeddings are aggregated using a permutation-invariant readout function to yield a molecular embedding that is mapped to target properties through downstream networks. The primary motivation for using GNNs is their ability to model molecular structure directly and provide atomistic interpretability. Node and edge representations capture subtle variations in heteroatom placement, aromatic substitution, and charge patterning not easily expressed with coarse descriptors. Because molecular embeddings are constructed from atom-level features, model outputs can be decomposed into contributions from individual atoms and functional groups, yielding chemically interpretable insights. GNNs typically outperform classical models on large datasets, but condensate partitioning data here are relatively small. To mitigate this, we use pretraining to initialize the graph encoder on a larger collection of molecules with related physicochemical properties before fine-tuning on condensate partitioning data, improving data efficiency while preserving interpretability.

For pretraining, molecular structures are represented as graphs from SMILES strings using Chemprop featurization, with atoms as nodes and bonds as directed edges. The GNN backbone employs bond-centered message-passing comprising five layers with 500 hidden units and SiLU activations. Node embeddings are updated through bond-conditioned transformations and aggregated via mean pooling to yield 500-dimensional molecular representations. The encoder was pretrained on 150,000 small molecules from ChEMBL [39], optimized to jointly predict logP, logS, and CIlogS using multitask regression, allowing the network to learn representations sensitive to hydrophobicity and solvation. **Fig. 2** shows the pretraining scheme.

### Regression model for small molecule condensate partitioning

For the regression task, the module contains four shared fully connected layers (500 hidden units, SiLU activation, dropout 0.08) that act as a common transformation for all downstream tasks. This shared representation is then branched into four independent task specific heads corresponding to the SUMOSIM, cGAS DNA, Dhh1, and SH3PRM systems. Each task head includes two fully connected layers with 500 hidden units that adapt the shared embedding to the respective task distribution. From these layers, each head produces four outputs: the evidential mean (*γ*) and shape parameters (*α, β*) defining a Normal Inverse Gamma (NIG) distribution for the *LogPC* target, and an auxiliary scalar predicting *QPlogKhsa*. The inclusion of *QPlogKhsa* as an auxiliary task was motivated by its moderate correlation with *LogPC* since it captures general hydrophobic and protein binding tendencies without being so highly correlated that it dominates the learning objective. The auxiliary supervision stabilizes shared representation learning and mitigates negative transfer between the condensate specific tasks:

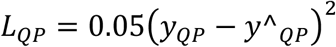

In parallel, each task possesses an independent lightweight sub network dedicated to the evidential parameter *v*. This pathway is separate from both the shared and task specific layers and takes the concatenated molecular embedding and auxiliary features as input. It directly outputs a single scalar v without sharing parameters with the *γ, α, β*, or *QPlogKhsa* branches. This design allows *v* to represent epistemic uncertainty independently of the predictive mean. Biases for *v* are initialized between 0.7 and 0.9 across tasks, while *α* and *β* are initialized to positive values ensuring numerical stability. Outputs from the four task heads and their corresponding *v* branches are concatenated into a 20-dimensional output vector ordered as SUMOSIM, cGAS-DNA, Dhh1, SH3PRM. For each molecule, the model thus predicts (*γ, v, α, β, QPlogKhsa*) for every condensate system.

The Normal Inverse Gamma (NIG) prior is defined as:

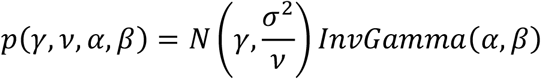

which implies a student t predictive distribution for the standardized *LogPC* target y:

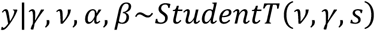

where 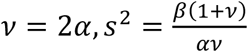

The per sample negative log likelihood loss used for training is:

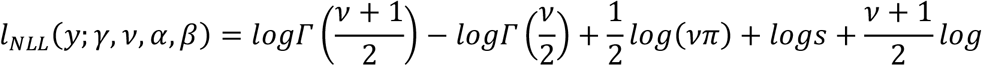

The predictive variance decomposes into aleatoric and epistemic terms:

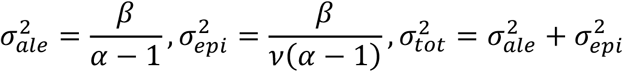

The total per-task loss combines these components with regularization:

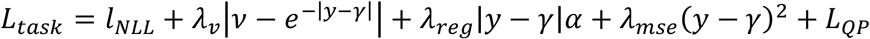

and the multitask batch loss is averaged across condensates:

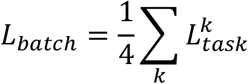

Training targets are standardized by subtracting the mean and dividing by the standard deviation computed on the training set for each property. Predictions are later transformed back to the original scale for evaluation. Training uses the AdamW optimizer (learning rate 7×10^−5^, weight decay 1×10^−6^, batch size 128) with cosine learning rate decay over 500 epochs. Regularization weight λ_reg_ is linearly annealed from 5×10^−4^ to 2×10^−3^ during the first 100 epochs, and early stopping monitors the mean validation R^2^ across all condensate systems with patience of 60 epochs. The model architecture is as shown in **Fig. 2**.

### SHAP Analysis

We use SHapley Additive exPlanations (SHAP) to interpret how the trained graph neural network attributes small molecule partitioning predictions to individual atoms. SHAP is a game theoretic attribution framework that assigns each feature a contribution value defined as its average marginal effect on the model output across all possible feature subsets. In our setting, features correspond to atoms in the molecular graph, and the model output of interest is the predicted logPC for a given condensate. To compute atom level SHAP values, we evaluate the model under systematic atom masking. For each molecule, we construct the Chemprop BatchMolGraph representation and define a binary mask vector with one entry per atom. Applying a mask sets the masked atom feature vector to zero and simultaneously attenuates edge features for any bond incident to a masked atom by multiplying the corresponding bond feature vector by the product of the source and destination atom masks. This preserves the molecular graph topology while removing information associated with selected atoms and their bonds. For each condensate, we compute SHAP values using a permutation based SHAP estimator, which approximates Shapley values by sampling random permutations of atom inclusion order and measuring changes in the predicted logPC as atoms are added back to the masked baseline. Unless otherwise specified, the baseline corresponds to the fully masked graph in which all atom features are zeroed. Because our uncertainty model outputs multiple quantities per condensate, we compute SHAP values with respect to the gamma head, which represents the model predicted logPC. For each molecule and condensate, we obtain an atom wise SHAP vector whose entries sum to the difference between the predicted logPC and the baseline prediction. We then aggregate atom level attributions across molecules and map each atom to a broad element category based on its atomic symbol, including C, N, O, S, P, F, Cl, Br, I, and Other. This broad grouping enables comparison of atomic contributions across chemically diverse molecules while retaining interpretability at the level of elemental functionality. For each condensate, we export per atom SHAP values and associated metadata, and we visualize the distribution of SHAP contributions for each element group using violin plots.

## Acknowledgements

This work was supported by the National Institute of General Medical Sciences of the National Institute of Health (R35GM153388). We gratefully acknowledge the Texas A&M High Performance Research Computing for providing the computational resources for this work.

## Notes

### Competing Interest Statement

The authors have declared no competing interest.

