## Supplementary Information for "Learning molecular determinants of selective small-molecule partitioning across biomolecular condensates"

### Selection of feature set

For feature selection, we systematically evaluated multiple molecular representation strategies to identify descriptor spaces that most effectively structured chemical space with respect to partitioning behavior. For each candidate feature set, molecular descriptors were computed and subsequently projected into two dimensions using t SNE to enable qualitative visualization of cluster organization. Representative projections are shown in Figs S3 to S10. To complement visual inspection with quantitative assessment, we computed clustering quality metrics including the Silhouette score, Davies Bouldin index, and Calinski Harabasz score for each representation. The Silhouette score evaluates how well molecules are matched to their assigned cluster relative to neighboring clusters. The Davies Bouldin index measures intra cluster compactness relative to inter cluster separation, with lower values indicating better separation. The Calinski Harabasz score quantifies the ratio of between cluster dispersion to within cluster dispersion, where higher values reflect more distinct clustering structure. Based on the combined evaluation of these metrics, we selected the top three performing feature sets for downstream modelling for the baseline model. The full set of quantitative scores is reported in Table S1. This procedure ensured that the chosen representations exhibited both strong structural organization in chemical space and favorable quantitative clustering characteristics prior to model development.

### Physicochemical and Structural Profiles of Unique Condensate Partitioners and Excluders

To further characterize condensate selectivity within the ChEMBL dataset, we analyzed the physicochemical and structural properties of molecules belonging to two classes highlighted in Fig S19. The first class consists of molecules that partition into exactly one condensate, illustrated here for selective partitioning into SUMOSIM, cGAS DNA, and Dhh1. The second class consists of molecules that are excluded from exactly one condensate, illustrated for exclusion from cGAS DNA, Dhh1, and SH3PRM. For these six subgroups we calculated a broader set of molecular descriptors and structural motifs. Fig S19 presents the subset of molecular properties that show the most pronounced differences across the subgroups, including fraction of sp3 hybridized carbon atoms, heavy atom count, heteroatom count, and topological polar surface area (TPSA), together with the prevalence of selected functional motifs. Functional group prevalence is reported as the percentage of molecules in each subgroup containing the indicated functionality.

The property distributions for molecules that partition into exactly one condensate are shown in Fig S19 a. Clear differences are observed across the three subgroups, indicating that selective entry into a single condensate is associated with distinct molecular property profiles. Molecules partitioning into Dhh1 tend to occupy lower ranges of heavy atom count, heteroatom count, and TPSA, consistent with a comparatively compact and less polar chemical space. In contrast, molecules partitioning into cGAS DNA extend toward higher fractions of sp3 carbon and broader TPSA values, indicating a more saturated and more polar population. Molecules partitioning into SUMOSIM fall between these two regimes across several properties but remain distinguishable in their overall distribution patterns. These results indicate that selective partitioning into a single condensate corresponds to structured variation in molecular size, polarity, heteroatom content, and degree of saturation.

The distributions for molecules excluded from exactly one condensate are shown in Fig S19 b and also display pronounced subgroup structure. Molecules excluded from Dhh1 shift toward higher fractions of sp3 carbon, larger heavy atom counts, increased heteroatom counts, and substantially higher TPSA, describing a population that is generally larger, more saturated, and more polar. By comparison, molecules excluded from SH3PRM cluster at lower heavy atom counts, lower heteroatom counts, and lower TPSA, defining a more compact and less polar subset. Molecules excluded from cGAS DNA occupy an intermediate profile, with broader distributions than the SH3PRM excluded class but generally lower polarity and molecular size than the Dhh1 excluded class. These patterns indicate that exclusion from a single condensate is also chemically organized and can be resolved through coordinated variation in molecular property space.

Functional group analysis further clarifies these subgroup distinctions. As shown in Fig S19 c, molecules that partition into exactly one condensate differ strongly in motif composition. Compounds partitioning into SUMOSIM exhibit high prevalence of amide, alcohol, ester, and tertiary amine groups. Molecules partitioning into cGAS DNA are enriched in tertiary amines together with alcohol containing functionalities. In contrast, Dhh1 partitioning molecules show increased prevalence of heteroaromatic nitrogen containing scaffolds, halogen substitution, and amide motifs. The patterns among molecules excluded from exactly one condensate are shown in Fig S19 d. Molecules excluded from Dhh1 are dominated by alcohol and ester containing compounds, whereas molecules excluded from SH3PRM show strong enrichment of tertiary amines, heteroaromatic nitrogens, and amide groups. Molecules excluded from cGAS DNA retain substantial amide prevalence together with contributions from heteroaromatic nitrogen, ester, and carboxylic acid functionalities.

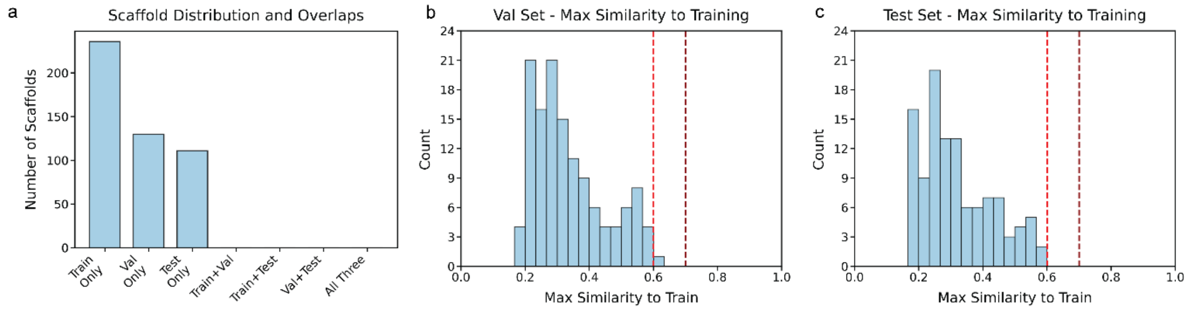


*Fig S1. Scaffold split quality checks. Panel a shows the number of unique Bemis Murcko scaffolds that appear only in the training set, only in the validation set, only in the test set, or are shared across splits. No shared scaffolds are observed, consistent with a strict scaffold-based split. Panels b and c show the distribution of the maximum fingerprint similarity of each validation molecule (b) or test molecule (c) to any molecule in the training set. The vertical dashed red lines indicate reference similarity cutoffs at 0.6 and 0.7 for identifying close analogs.*

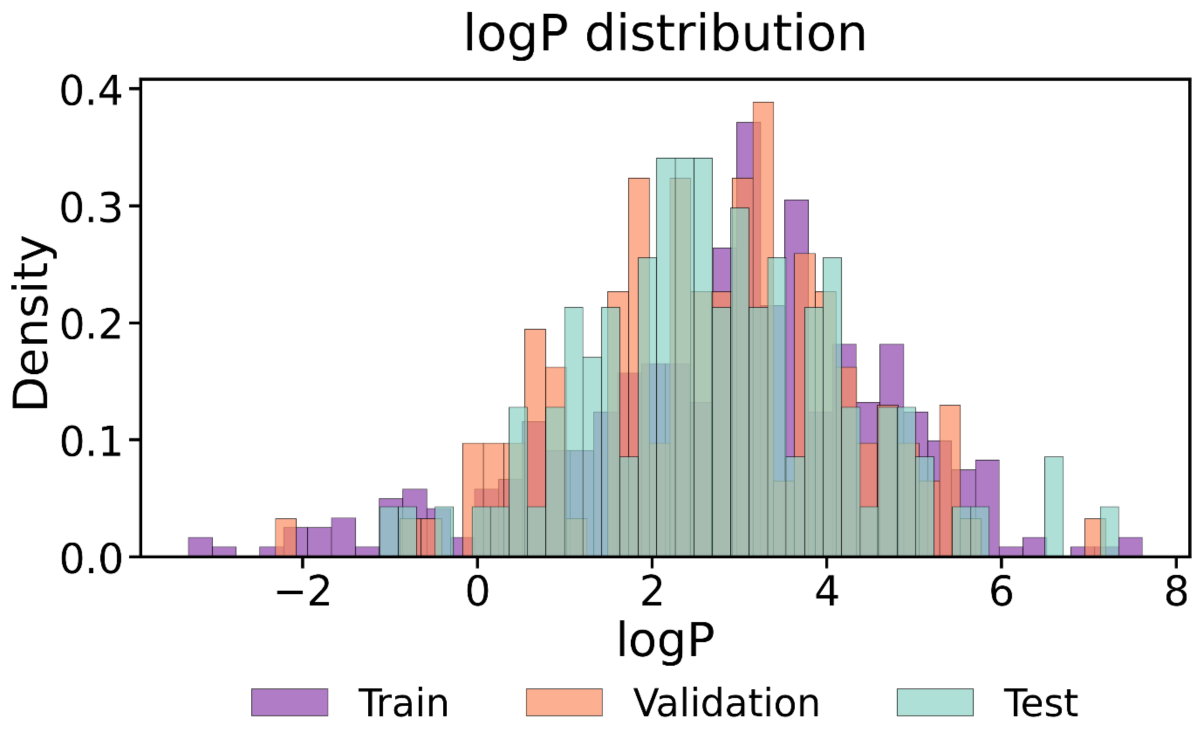


*Fig S2. Distribution of logP values across the training, validation, and test sets. We split the dataset by molecular structure to prevent leakage of chemically similar compounds between training and evaluation sets. This ensures that model performance reflects true generalization across chemical space rather than interpolation between closely related molecules. Lipophilicity logP was not used as a splitting criterion because it is a bounded scalar property shared by many structurally diverse molecules. As long as the logP distribution of the evaluation set lies within the support of the training set the model is evaluated under in distribution conditions with respect to lipophilicity while remaining out of distribution in molecular structure. This allows the assessment to focus on extrapolation across chemotypes rather than conflating structural generalization with property range extrapolation. The resulting setup reflects the intended use case where novel chemical structures are encountered within an established and physically reasonable lipophilicity range.*

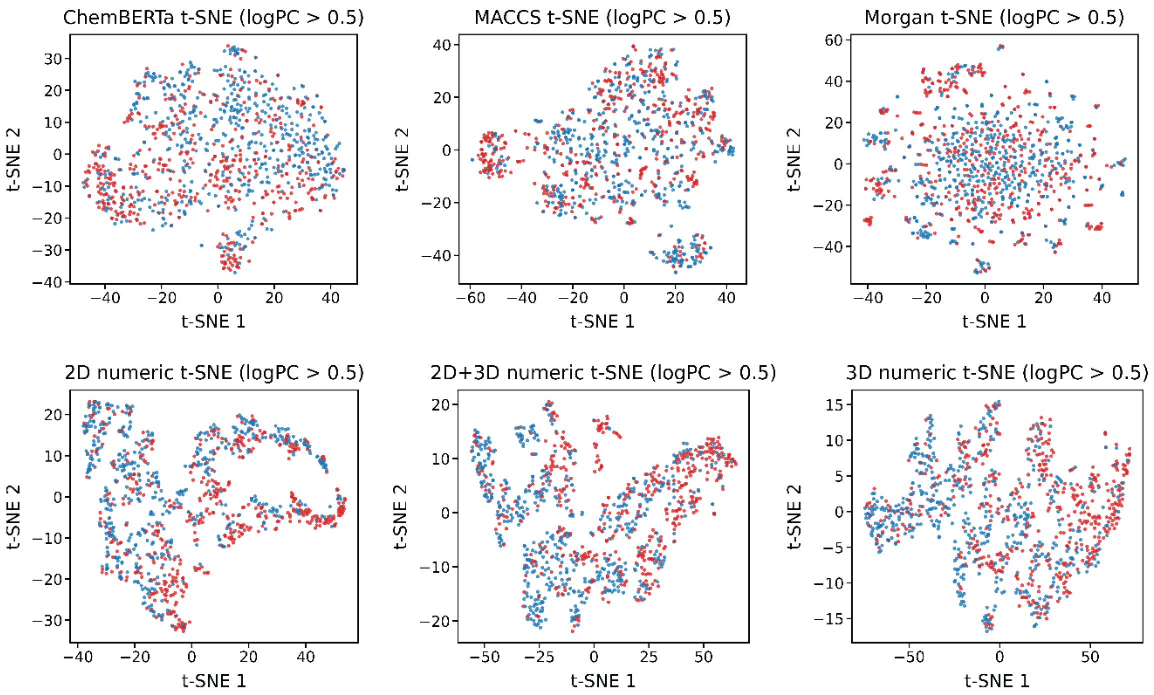


*Fig S3. t-SNE visualization of molecular representations for compounds associated with the cGAS DNA condensate. Each panel shows a two-dimensional projection of a different descriptor space including ChemBERTa embeddings, MACCS keys (167 bit), Morgan fingerprints (radius 2, 2048 bits), and numerical feature sets derived from 2D descriptors, 3D descriptors, and their combination. Points are colored according to partitioning behavior. The plots allow qualitative comparison of how different featurization strategies organize chemical space and how effectively they separate partitioning from non-partitioning molecules.*

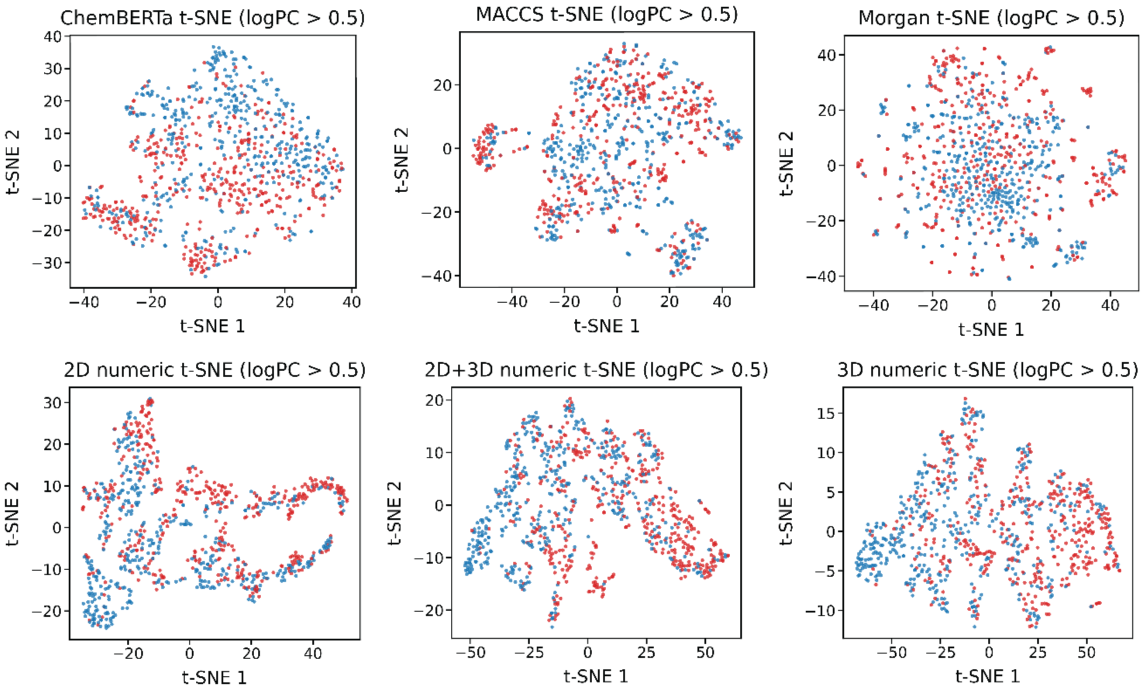


*Fig S4. t-SNE visualization of molecular representations for compounds associated with the Dhh1 condensate. Each panel shows a two-dimensional projection of a different descriptor space including ChemBERTa embeddings, MACCS keys (167 bit), Morgan fingerprints (radius 2, 2048 bits), and numerical feature sets derived from 2D descriptors, 3D descriptors, and their combination. Points are colored according to partitioning behavior. The plots allow qualitative comparison of how different featurization strategies organize chemical space and how effectively they separate partitioning from non-partitioning molecules.*

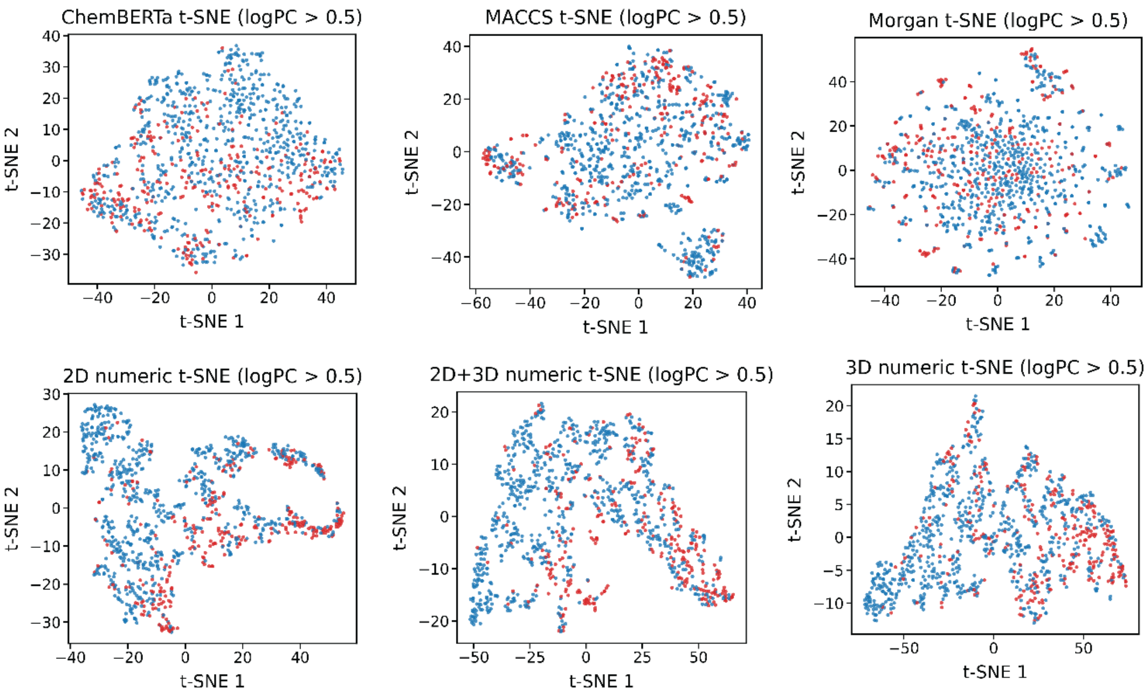


*Fig S5. t-SNE visualization of molecular representations for compounds associated with the SH3PRM condensate. Each panel shows a two-dimensional projection of a different descriptor space including ChemBERTa embeddings, MACCS keys (167 bit), Morgan fingerprints (radius 2, 2048 bits), and numerical feature sets derived from 2D descriptors, 3D descriptors, and their combination. Points are colored according to partitioning behavior. The plots allow qualitative comparison of how different featurization strategies organize chemical space and how effectively they separate partitioning from non-partitioning molecules.*

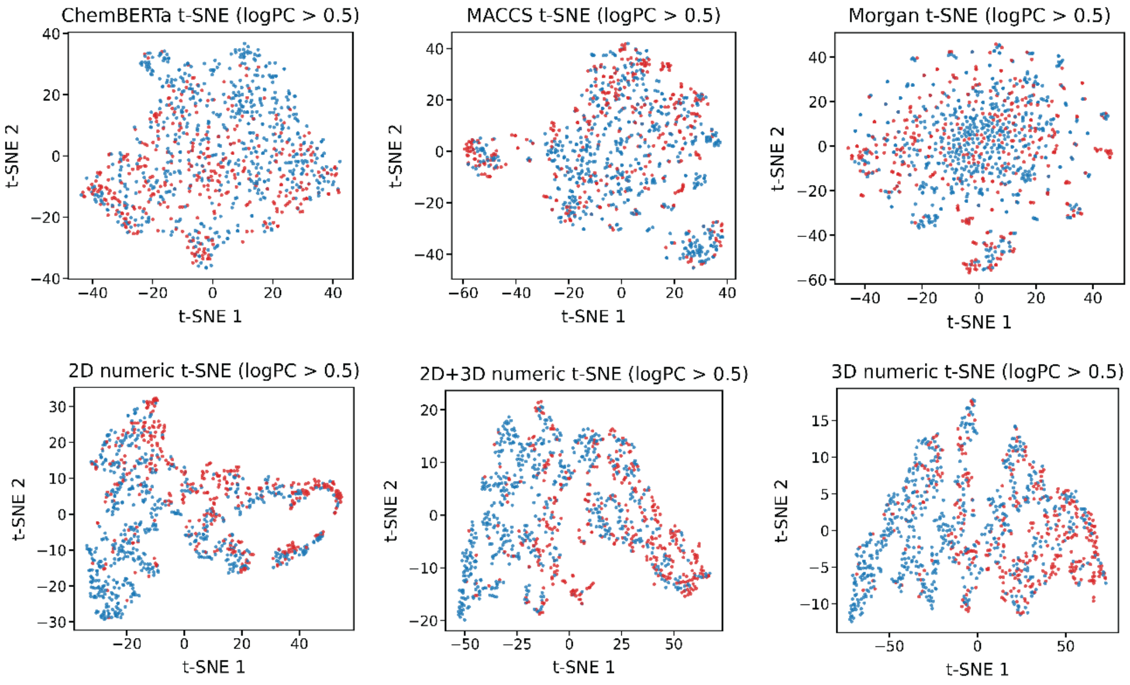


*Fig S6. t-SNE visualization of molecular representations for compounds associated with the SUMOSIM condensate. Each panel shows a two-dimensional projection of a different descriptor space including ChemBERTa embeddings, MACCS keys (167 bit), Morgan fingerprints (radius 2, 2048 bits), and numerical feature sets derived from 2D descriptors, 3D descriptors, and their combination. Points are colored according to partitioning behavior. The plots allow qualitative comparison of how different featurization strategies organize chemical space and how effectively they separate partitioning from non-partitioning molecules.*

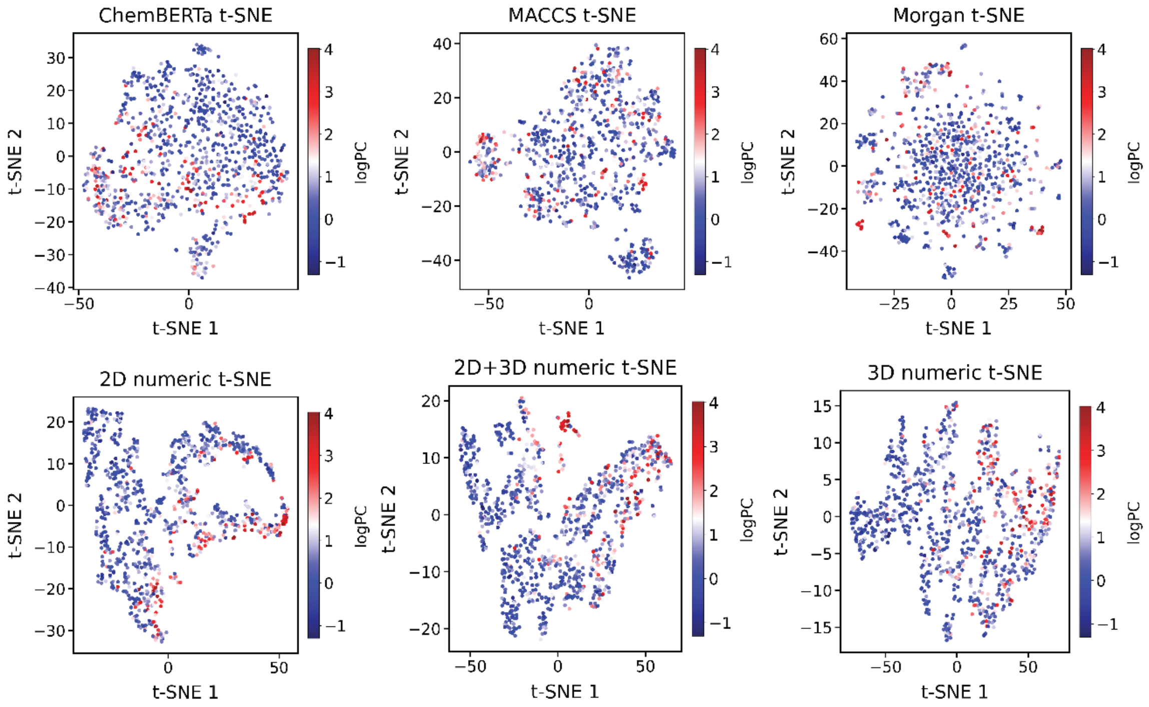


*Fig S7. t-SNE visualization of molecular representations for the cGAS DNA condensate regression dataset. Each panel shows a two-dimensional projection of a different descriptor space including ChemBERTa embeddings, MACCS keys (167 bit), Morgan fingerprints (radius 2, 2048 bits), and numerical feature sets derived from 2D descriptors, 3D descriptors, and their combination. Points are colored by the measured partition coefficient values used as the regression target. The projections allow qualitative assessment of how different featurization strategies organize chemical space and whether the target property exhibits structured gradients within each representation.*

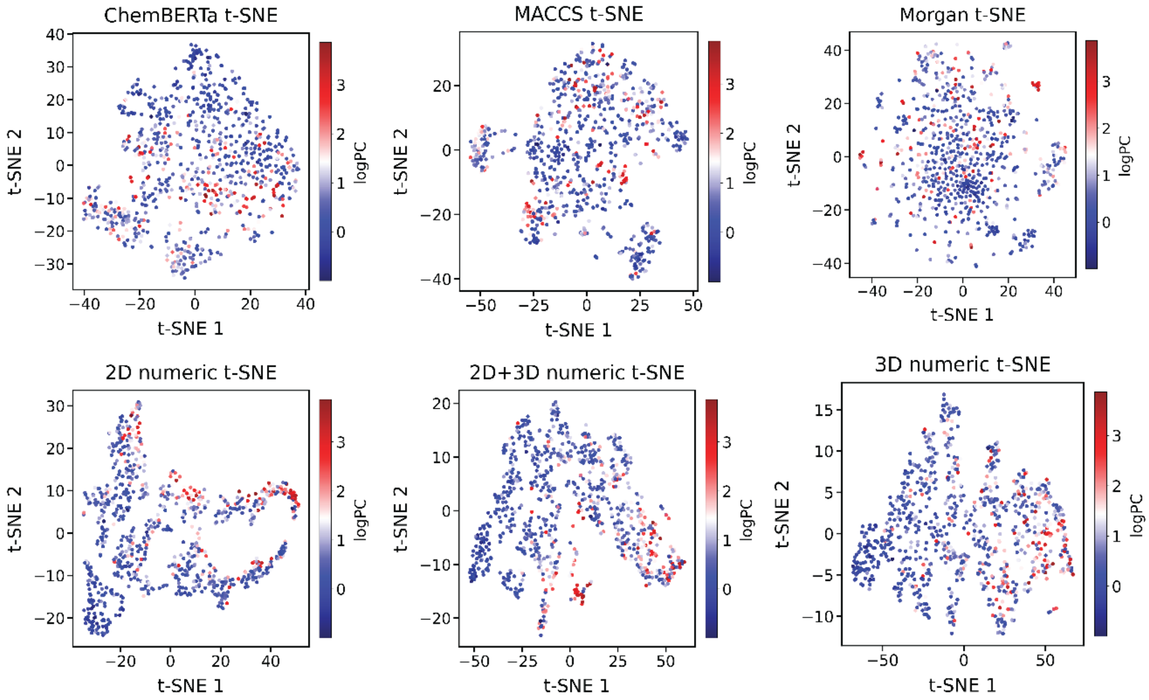


*Fig S8. t-SNE visualization of molecular representations for the Dhh1 condensate regression dataset. Each panel shows a two-dimensional projection of a different descriptor space including ChemBERTa embeddings, MACCS keys (167 bit), Morgan fingerprints (radius 2, 2048 bits), and numerical feature sets derived from 2D descriptors, 3D descriptors, and their combination. Points are colored by the measured partition coefficient values used as the regression target. The projections allow qualitative assessment of how different featurization strategies organize chemical space and whether the target property exhibits structured gradients within each representation.*

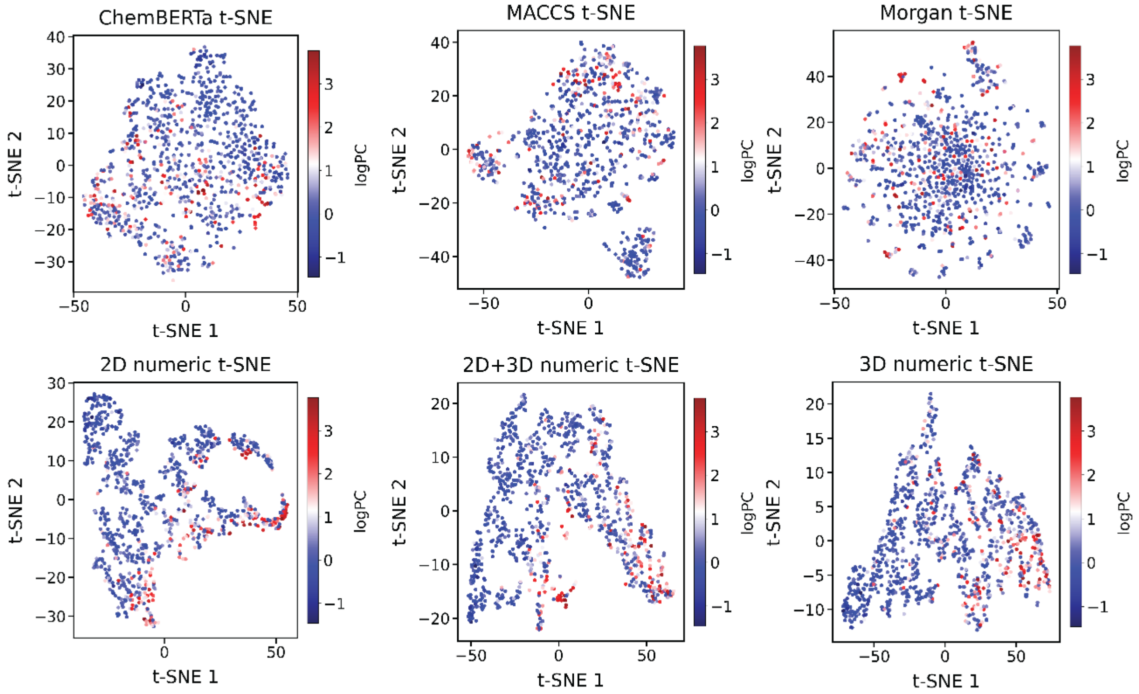


*Fig S9. t-SNE visualization of molecular representations for the SH3PRM condensate regression dataset. Each panel shows a two-dimensional projection of a different descriptor space including ChemBERTa embeddings, MACCS keys (167 bit), Morgan fingerprints (radius 2, 2048 bits), and numerical feature sets derived from 2D descriptors, 3D descriptors, and their combination.. Points are colored by the measured partition coefficient values used as the regression target. The projections allow qualitative assessment of how different featurization strategies organize chemical space and whether the target property exhibits structured gradients within each representation.*

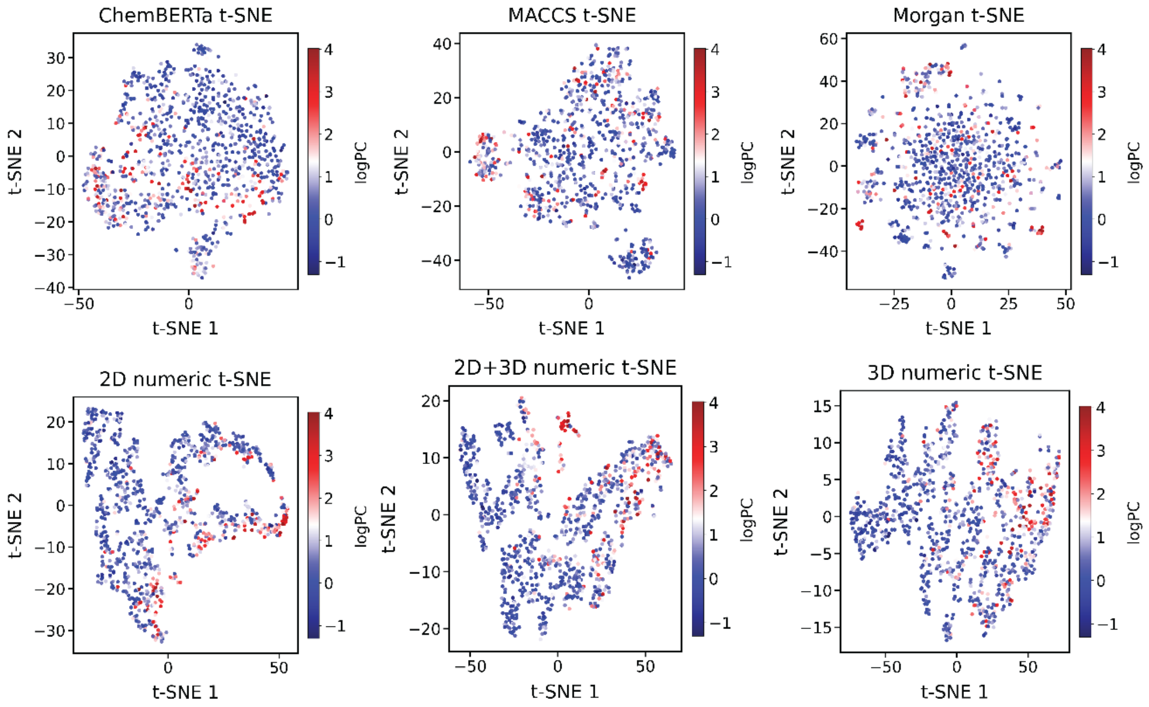


*Fig S10. t-SNE visualization of molecular representations for the SUMOSIM condensate regression dataset. Each panel shows a two-dimensional projection of a different descriptor space including ChemBERTa embeddings, MACCS keys (167 bit), Morgan fingerprints (radius 2, 2048 bits), and numerical feature sets derived from 2D descriptors, 3D descriptors, and their combination. Points are colored by the measured partition coefficient values used as the regression target. The projections allow qualitative assessment of how different featurization strategies organize chemical space and whether the target property exhibits structured gradients within each representation.*

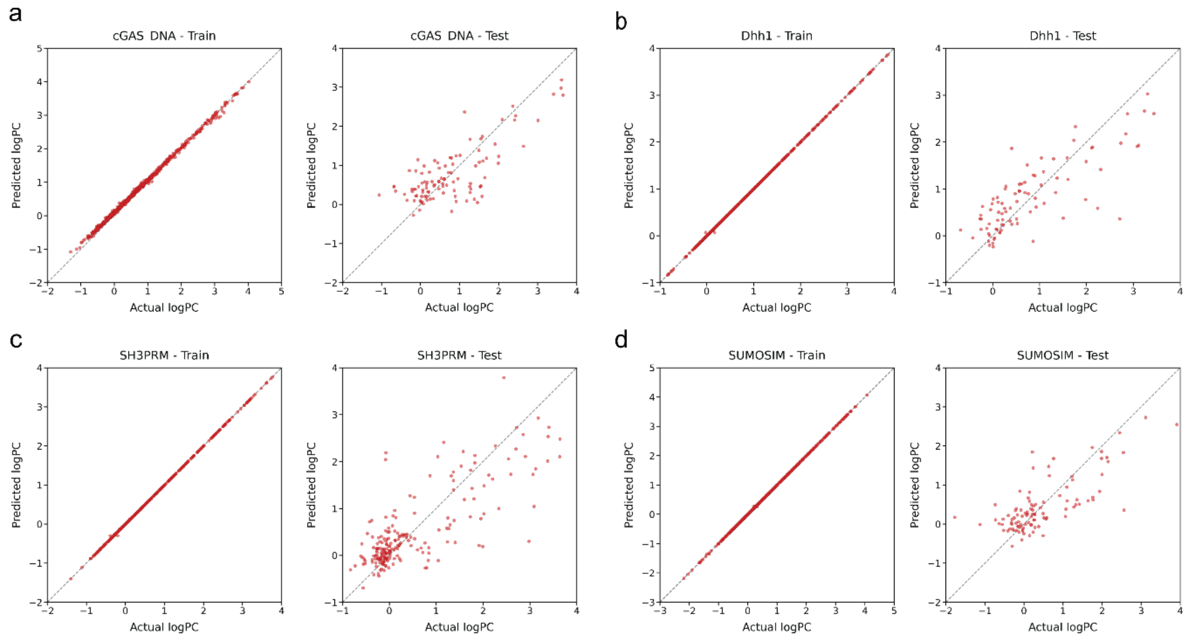


*Fig S11. Parity plots for baseline model performance across condensates. Panels a to d show predicted versus experimental logPC for cGAS DNA, Dhh1, SH3PRM, and SUMOSIM, respectively, with training results on the left and test results on the right in each panel. The gray dashed diagonal indicates the ideal y = x agreement line.*

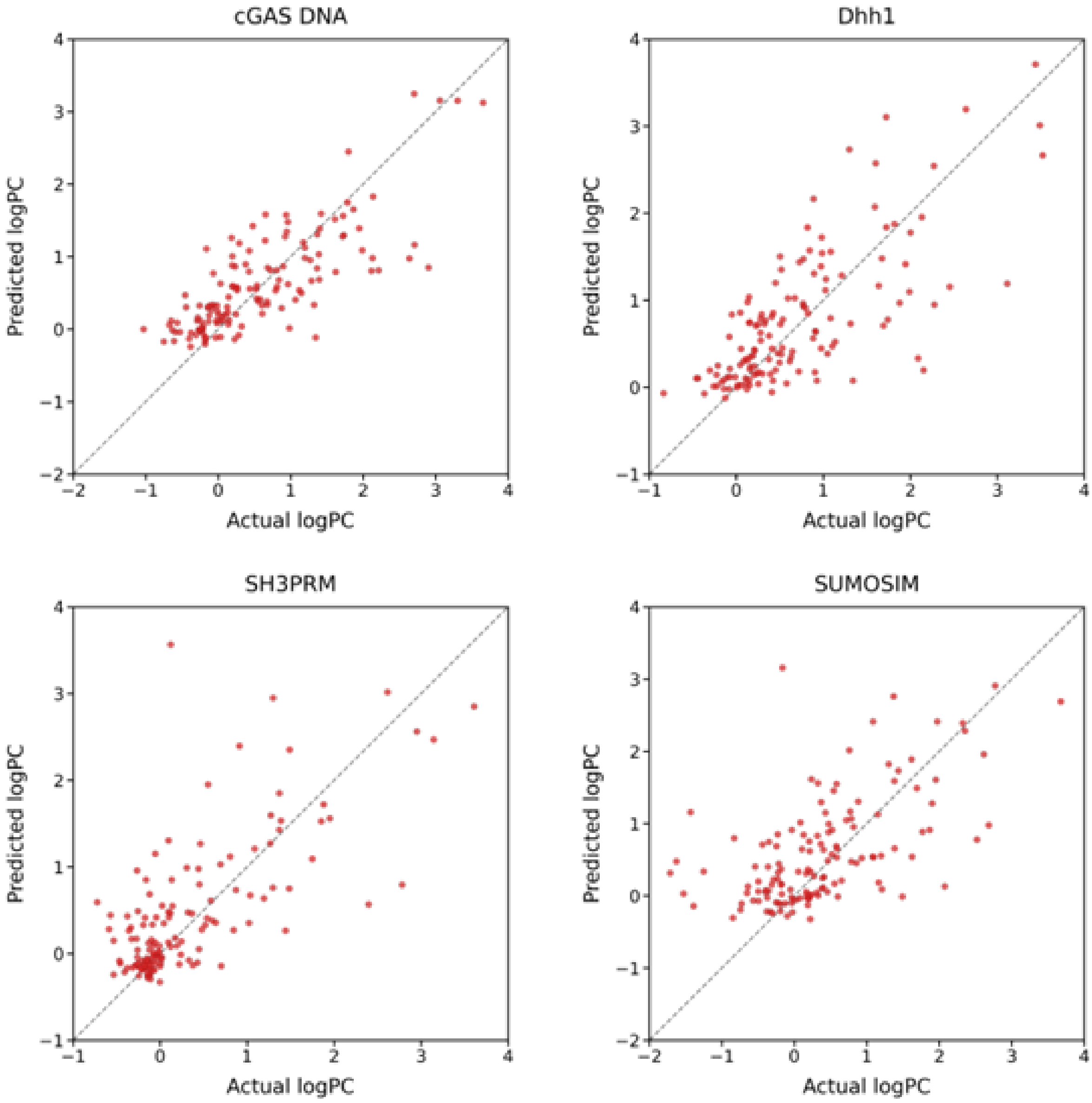


*Fig S12. Base GNN performance for condensate partitioning on the test sets.*

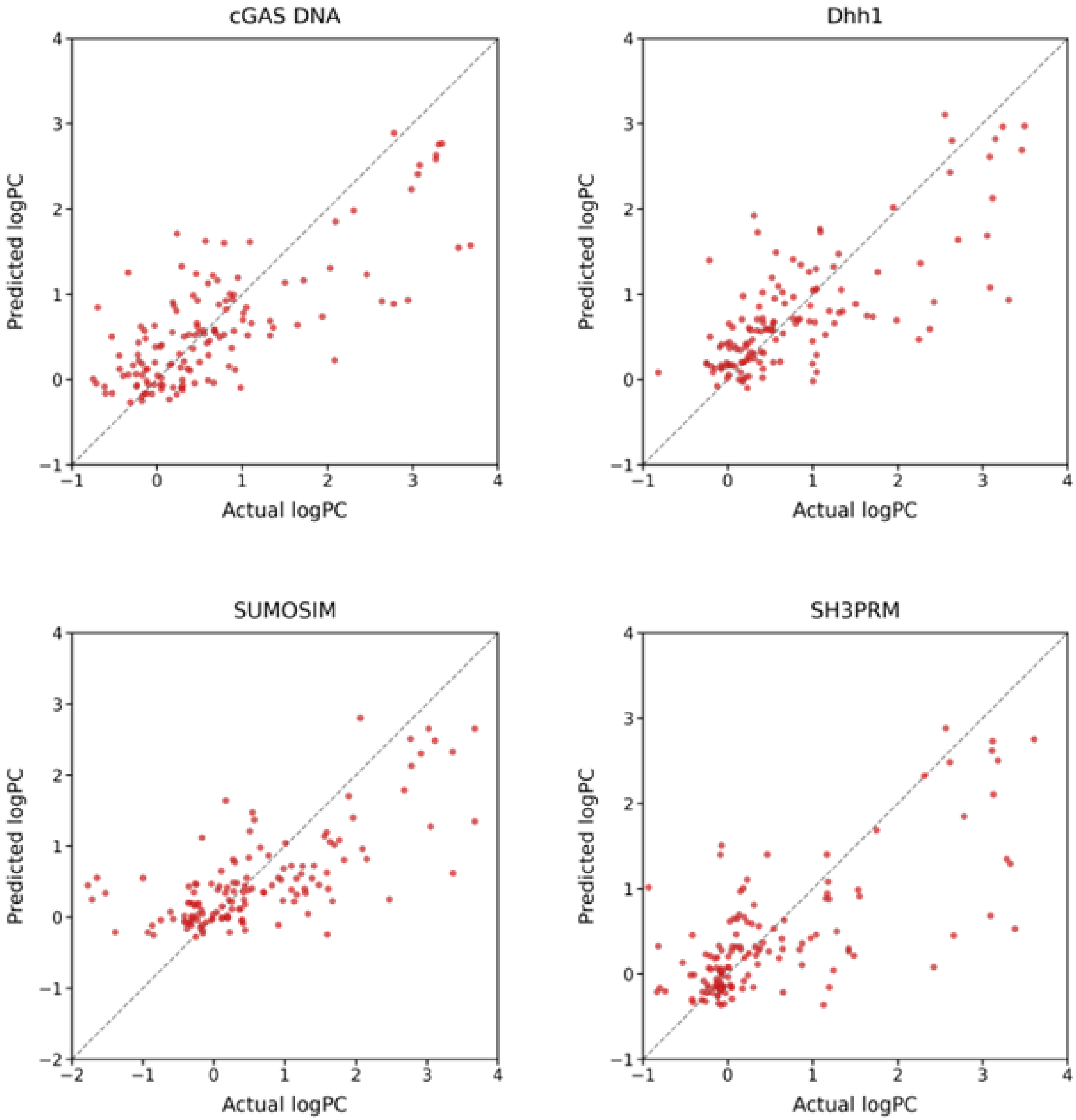


*Fig S13. Pre-trained GNN performance for condensate partitioning on the test sets.*

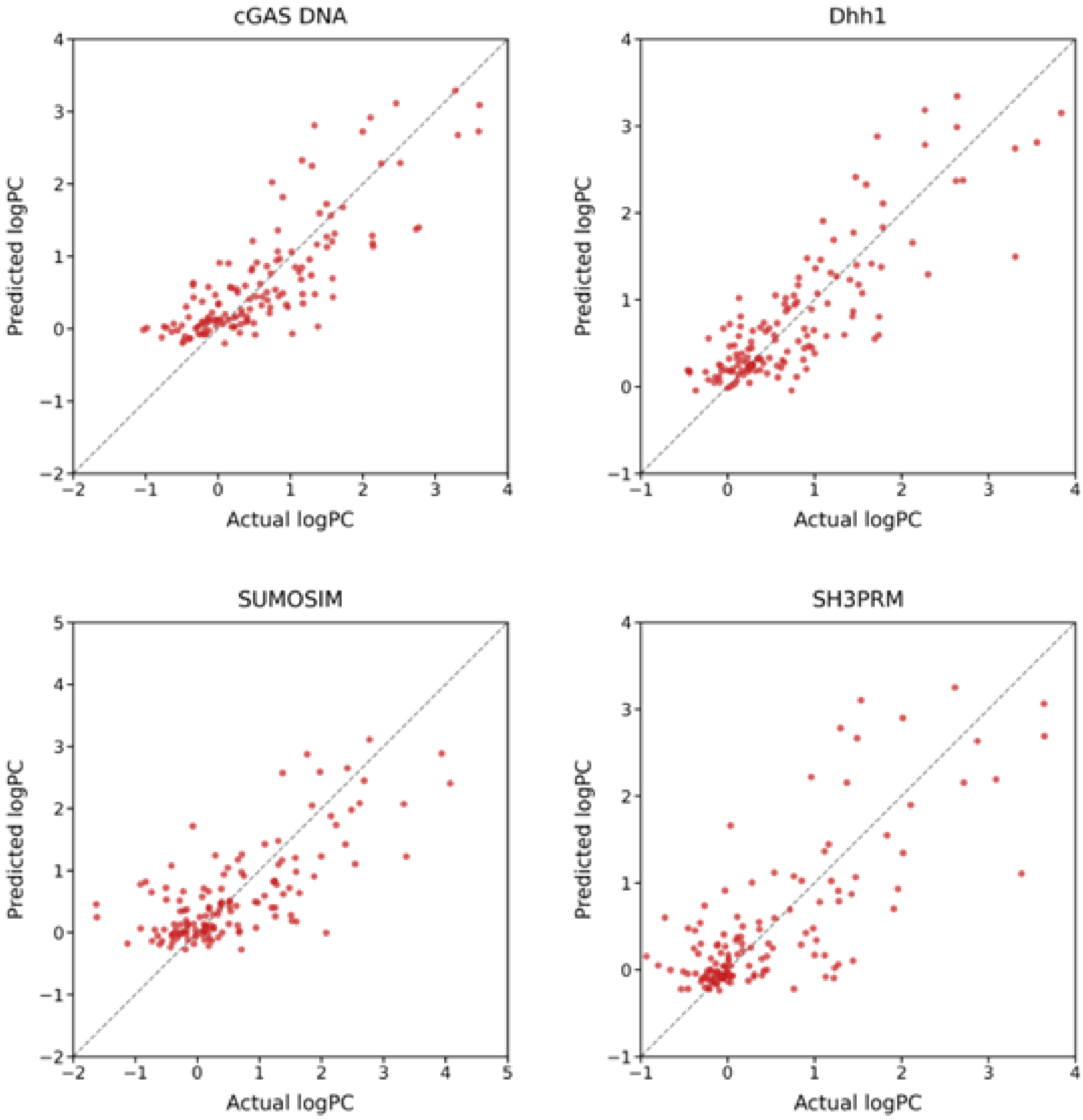


*Fig S14. Multitask GNN performance for condensate partitioning on the test sets.*

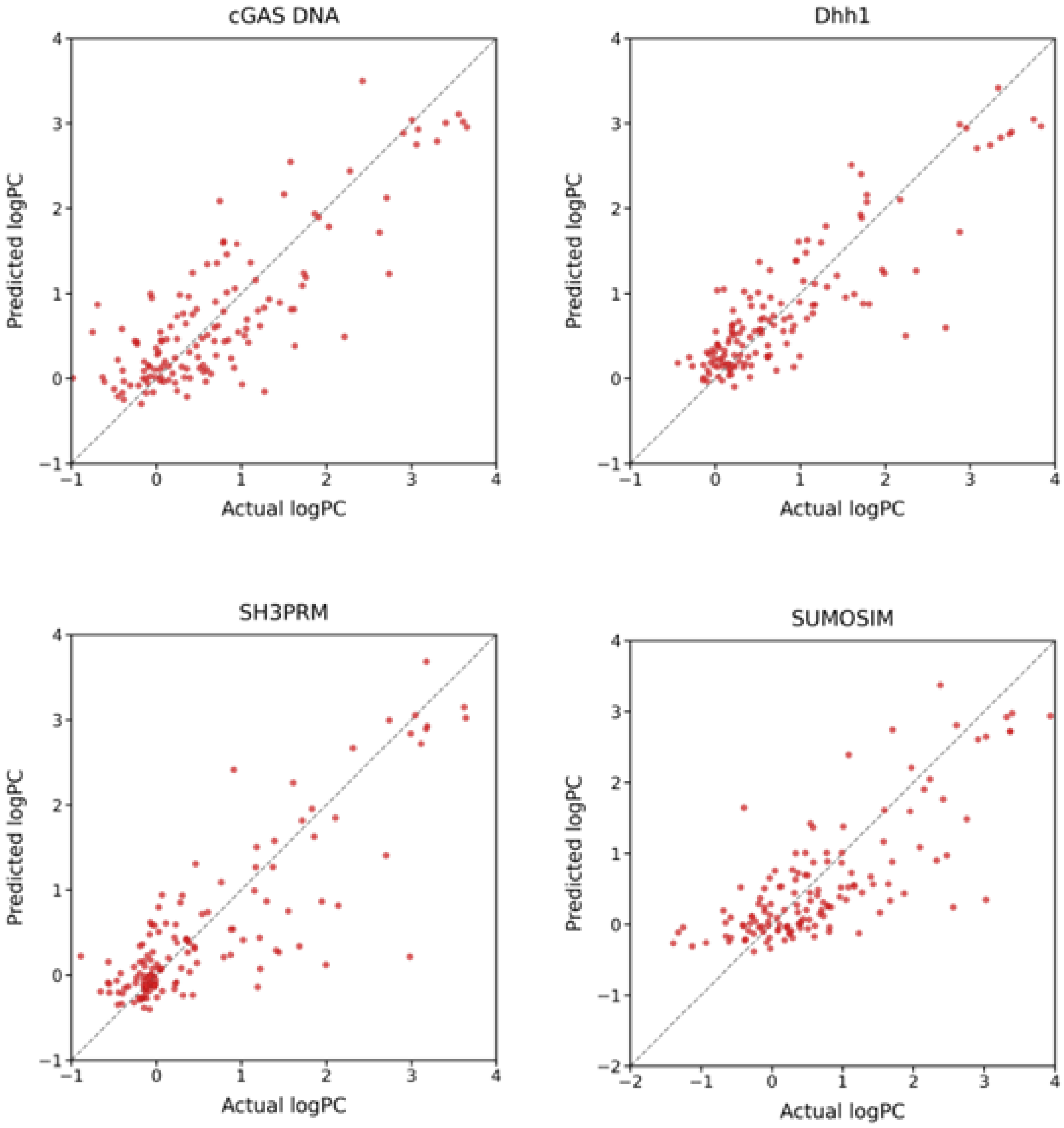


*Fig S15. Auxillary GNN performance for condensate partitioning on the test sets.*

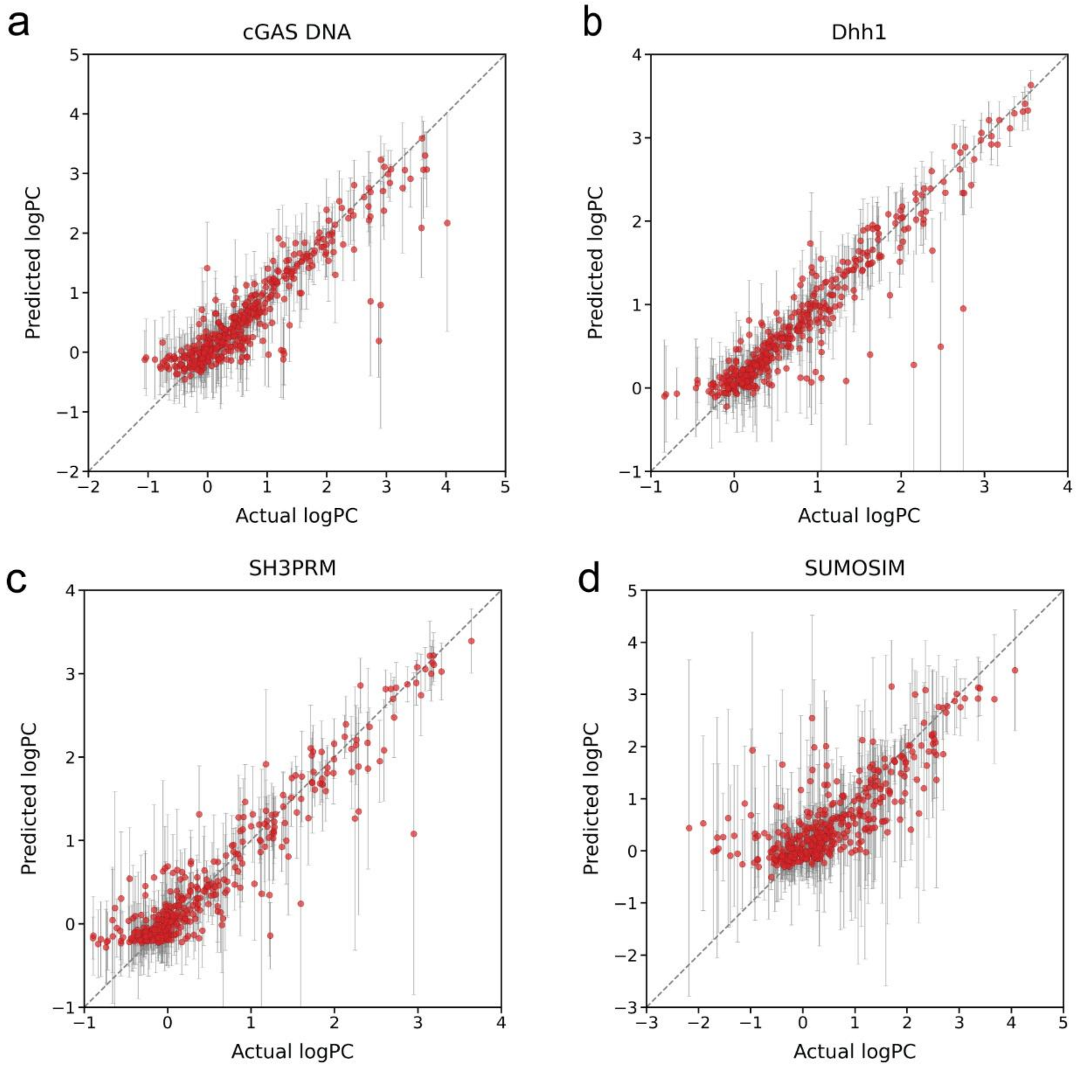


*Fig S16. Evidential regression performance for condensate partitioning on the training sets. Panels a to d show parity plots comparing predicted and experimental logPC for cGAS DNA, Dhh1, SH3PRM, and SUMOSIM, respectively. Each point corresponds to a single molecule and the vertical error bars indicate the model inferred predictive standard deviation for that molecule. The gray dashed diagonal indicates ideal agreement.*

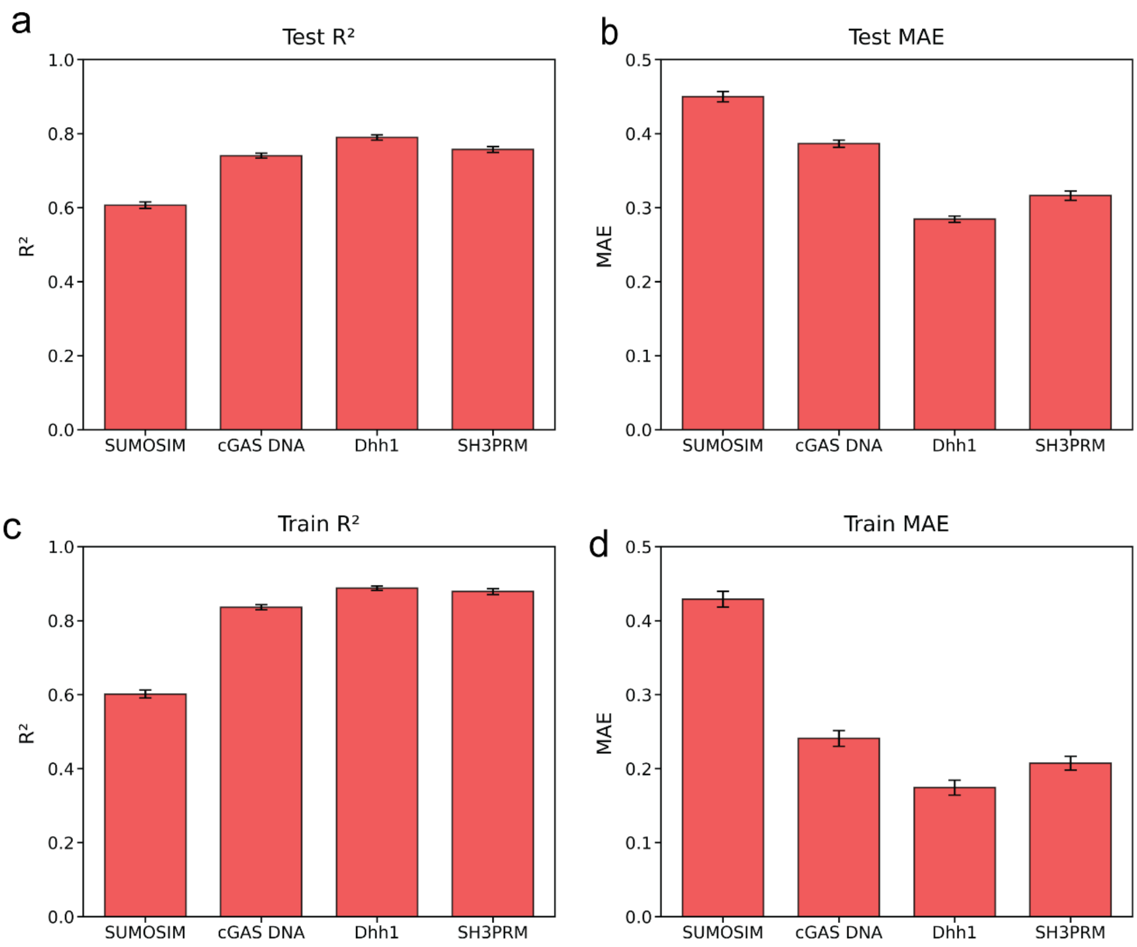


*Fig S17. Robustness across random initializations. Performance of the evidential regression models trained with 50 independent random initializations, reported as mean plus or minus standard error for each condensate. Panels a and b show test set $R^2$ and MAE, respectively, while panels c and d show the corresponding training set $R^2$ and MAE.*

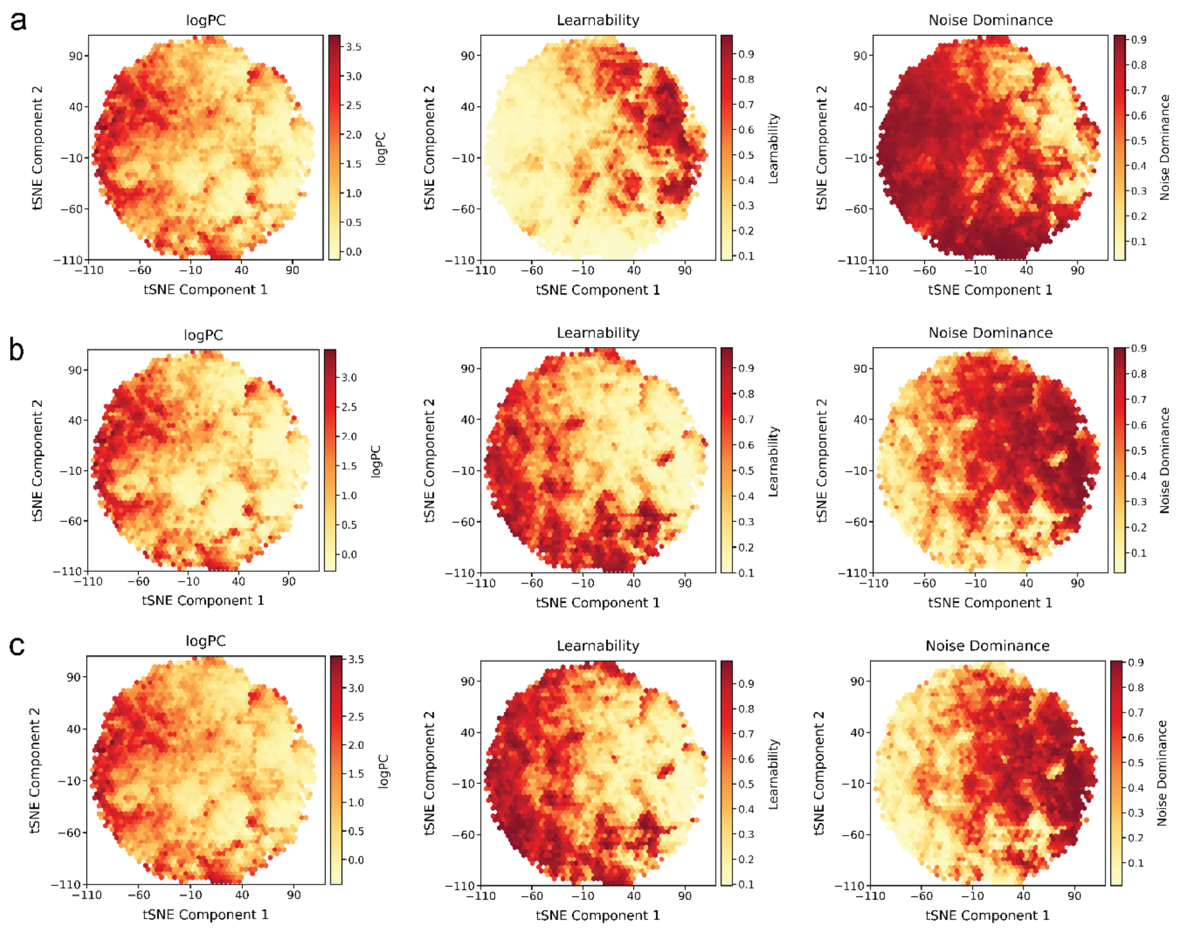


*Fig S18. Two-dimensional t-SNE maps of chemical space with hexagonal bin aggregation for three condensate systems. Rows a, b, and c correspond to Dhh1, SH3PRM, and SUMOSIM respectively. Within each row, the left panel shows the spatial distribution of logPC values, the middle panel shows learnability scores, and the right panel shows noise dominance.*

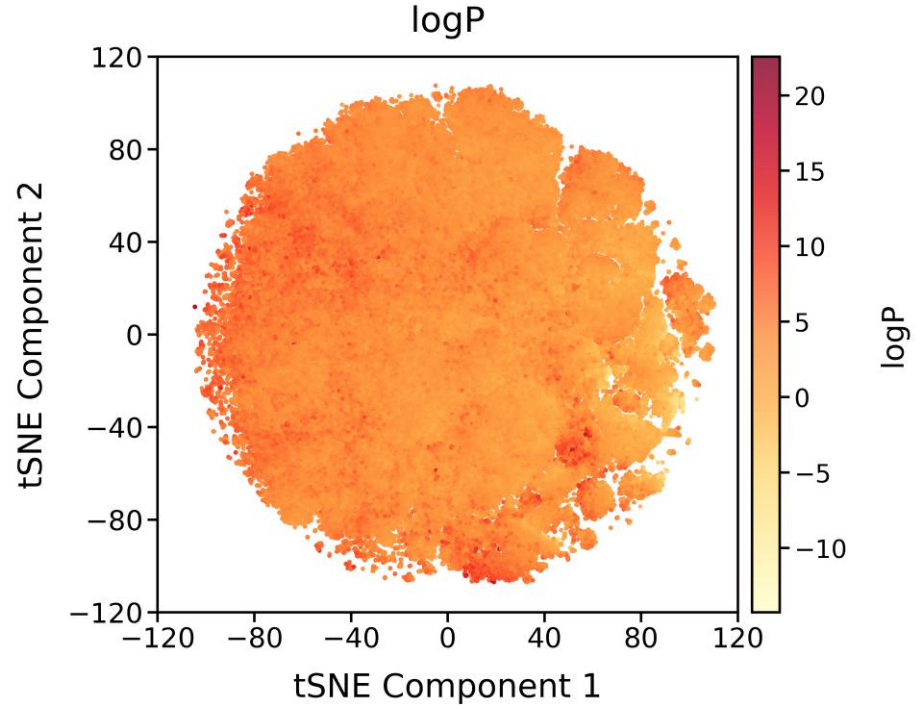


*Fig S19. The tSNE projections colored by logP show that lipophilicity is broadly distributed across the embedding space without clear spatial organization or clustering patterns. In contrast to the structured distribution observed for LogPC, the logP values appear diffuse across the projection, indicating that the representation is not primarily organized according to lipophilicity and suggesting that the LogPC structure does not arise from bias carried over from pretraining during fine tuning.*

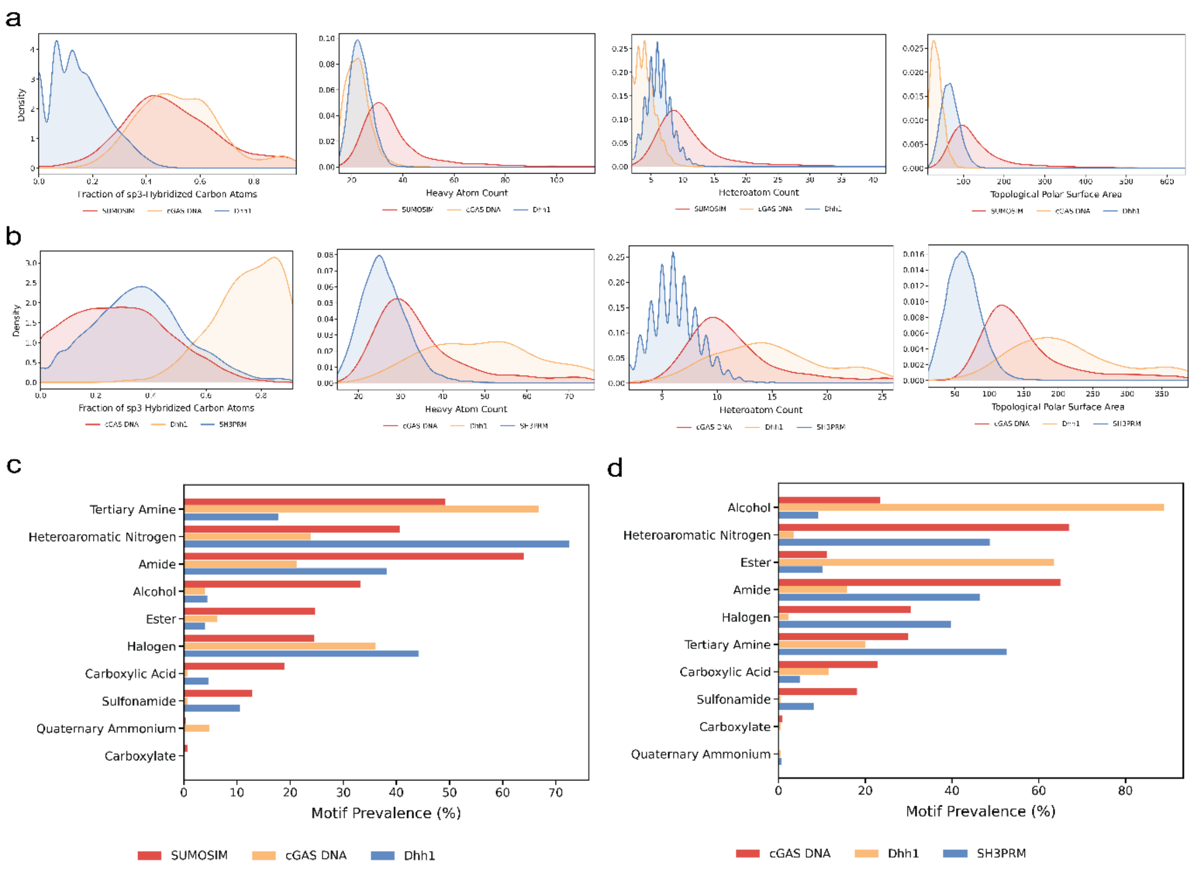


*Fig S20: (a) Kernel density estimates of the fraction of sp3-hybridized carbon atoms, heavy atom count, heteroatom count, and topological polar surface area (TPSA) for ChEMBL molecules that partition into exactly 1 condensate, grouped by the condensate entered (SUMOSIM, cGAS_DNA, and Dhh1). (b) Kernel density estimates of the fraction of sp3-hybridized carbon atoms, heavy atom count, heteroatom count, and topological polar surface area (TPSA) for ChEMBL molecules excluded from exactly 1 condensate, grouped by the condensate excluded (cGAS_DNA, Dhh1, and SH3PRM). (c) Prevalence of selected structural motifs for ChEMBL molecules that partition into exactly 1 condensate, grouped by the condensate entered (SUMOSIM, cGAS_DNA, and Dhh1). (d) Prevalence of selected structural motifs for ChEMBL molecules excluded from exactly 1 condensate, grouped by the condensate excluded (cGAS_DNA, Dhh1, and SH3PRM).*

*Table S1. Separability metrics for the various feature sets*

| Feature set | Silhouette score | Calinski harabasz | Davies bouldin |
|---|---|---|---|
| Chemberta | 0.058 | 51.89 | 3.79 |
| MAACS | 0.009 | 4.41 | 12.85 |
| Morgan fingerprints | 0.023 | 10.68 | 8.12 |
| 2d+3d features | 0.11 | 152.9 | 2.14 |
| 2d features | 0.057 | 53.46 | 3.66 |
| 3d features | 0.124 | 181.91 | 1.9 |

*Table S2. Tuned baseline model hyperparameters for each condensate including feature set and final values for all optimized training parameters.*

| Condensate | SUMOSIM | cGAS DNA | Dhh1 | SH3PRM |
|---|---|---|---|---|
| Feature_Set | qikprop | qikprop | all | all |
| Model | xgb | xgb | xgb | xgb |
| n_estimators | 1730 | 632 | 1788 | 2994 |
| learning_rate | 0.124317383 | 0.011682181 | 0.077589 | 0.001864 |
| max_depth | 11 | 9 | 3 | 10 |
| min_child_weight | 2.721786042 | 1.013633887 | 2.7251 | 3.889689 |
| subsample | 0.901254085 | 0.998850597 | 0.922243 | 0.991378 |
| colsample_bytree | 0.973426594 | 0.972566723 | 0.97834 | 0.773247 |
| reg_alpha | 2.43599E-08 | 1.05563E-06 | 2.44E-07 | 2.69E-08 |
| reg_lambda | 1.43622E-07 | 5.79767E-06 | 0.087875 | 0.051194 |
| gamma | 0.031585087 | 0.000389305 | 1.67E-07 | 0.002459 |

*Table S3. Model performance for the pre-training*

| Task | Train R2 | Test R2 | Train MAE | Test MAE |
|---|---|---|---|---|
| logP | 0.98 | 0.98 | 0.18 | 0.19 |
| logS | 0.97 | 0.97 | 0.23 | 0.24 |
| CIlogS | 0.98 | 0.98 | 0.3 | 0.31 |

*Table S4. Model performance for the various GNN variants*

| Condensate | Base GNN | | Pre-trained GNN | | Multitask GNN | | Auxiliary GNN | |
|---|---|---|---|---|---|---|---|---|
| | R2 | MAE | R2 | MAE | R2 | MAE | R2 | MAE |
| cGAS DNA | 0.43 | 0.49 | 0.58 | 0.48 | 0.66 | 0.44 | 0.68 | 0.44 |
| Dhh1 | 0.48 | 0.45 | 0.56 | 0.43 | 0.70 | 0.36 | 0.75 | 0.36 |
| SH3PRM | 0.40 | 0.40 | 0.51 | 0.46 | 0.56 | 0.42 | 0.70 | 0.37 |
| SUMOSIM | 0.25 | 0.58 | 0.48 | 0.59 | 0.52 | 0.55 | 0.57 | 0.54 |
